# Targeting the *XBP1* splice substrate redirects IRE1-mediated RNA processing without direct inhibition of IRE1

**DOI:** 10.64898/2026.11.26.769414

**Authors:** Anna Więch-Walów, Magdalena Gebert, Jakub Sławski, Paulina Czechowicz, Sylwia Bartoszewska, Diana Pelizzari-Raymundo, Wojciech Białek, Leszek Kalinowski, Liliana Schaefer, James F. Collawn, Eric Chevet, Rafal Bartoszewski

## Abstract

The endoplasmic reticulum (ER) stress sensor IRE1α catalyzes the unconventional splicing of X-box binding protein 1 (*XBP1*) mRNA together with the tRNA ligase RTCB, thus generating the transcription factor XBP1s, a central regulator of the unfolded protein response. Current approaches for suppressing XBP1 signaling primarily rely on inhibition of IRE1 kinase or RNase activities, thereby affecting multiple downstream outputs of the pathway. Here, we investigated whether direct targeting of the *XBP1* substrate could provide a more selective strategy for modulating *XBP1* splicing.

We designed a panel of antisense morpholino oligonucleotides directed against the *XBP1* splicing region and identified a morpholino, M2, that efficiently suppressed *XBP1s* production upon treatment with multiple ER stressors and in various cell lines. M2 treatment reduced XBP1s accumulation at both the transcript and protein levels without detectable effects on IRE1 phosphorylation or ATP conversion to ADP or RNase activity *in vitro*. In contrast to the selective IRE1 RNase inhibitor 4μ8C, M2 did not prevent ER stress-induced downregulation of the RIDD substrate *DGAT2*. Analysis of RNA turnover revealed increased persistence of RNA species containing the *XBP1u* region following M2 treatment, consistent with altered processing of the *XBP1* splice substrate rather than enhanced transcript degradation. Consistent with these observations, *in vitro* cleavage assays demonstrated that M2 altered the pattern of IRE1-mediated processing of *XBP1* RNA, resulting in accumulation of a cleavage intermediate while preserving detectable cleavage activity. Collectively, our findings demonstrate that selective targeting of the *XBP1* splice substrate is sufficient to suppress *XBP1s* production without direct inhibition of IRE1 and RIDD. These results establish substrate-directed modulation as a complementary strategy to enzyme-directed inhibition, and for the first time identify an additional level at which the IRE1–XBP1 pathway can be experimentally controlled to selectively dissociate RIDD outputs from that of *XBP1* splicing.

## INTRODUCTION

The unfolded protein response (UPR) is a conserved signaling network that maintains endoplasmic reticulum (ER) proteostasis under conditions of protein-folding stress. A central component of this response is inositol-requiring enzyme 1α (IRE1α), an ER-resident transmembrane protein that functions as both a kinase and endoribonuclease. Upon ER stress, IRE1 oligomerizes subsequently leading to activation of its RNase domain and initiating adaptive signaling pathways that promote restoration of ER homeostasis and cellular fitness [1, 2].

A major function of mammalian IRE1 signaling is the unconventional splicing of X-box binding protein 1 (*XBP1*) mRNA. Unlike canonical nuclear pre-mRNA splicing, this process occurs in the cytoplasm and involves site-specific cleavage of the unspliced *XBP1* transcript (*XBP1u*) at two conserved stem-loop structures flanking a 26-nucleotide intron. Ligation of the cleavage products by the tRNA ligase RTCB generates the spliced *XBP1s* transcript, which encodes a potent transcription factor controlling the expression of genes involved in protein folding, secretion, ER biogenesis and stress adaptation [1–3].

Efficient *XBP1* splicing requires not only recognition of specific RNA sequence and structural elements but also precise spatial organization of the substrate at ER-associated IRE1 signaling sites. Previous studies demonstrated that *XBP1u* mRNA is actively recruited to the ER membrane and that translational pausing encoded within the *XBP1u* transcript promotes efficient positioning of the substrate for IRE1-mediated cleavage. These findings established that unconventional *XBP1* splicing depends on specialized mechanisms governing both substrate recognition and localization [4, 5]. In addition to *XBP1* splicing, activated IRE1 mediates regulated IRE1-dependent decay (RIDD), a process involving endonucleolytic cleavage of a subset of RNA. Although both activities are catalyzed by the same RNase domain, accumulating evidence indicates that *XBP1* splicing and RIDD represent mechanistically distinct outputs of IRE1 signaling which are however connected since RIDD activity is increased in *XBP1* knock-out cells [6–9]. These processes differ in their substrate-recognition requirements and catalytic organization and can be experimentally uncoupled, suggesting that individual IRE1 outputs may be selectively manipulated without globally inhibiting IRE1 function [10–12].

Current approaches for modulating the IRE1-XBP1 pathway primarily target the catalytic activity of IRE1 itself, therefore preventing thus far to alter *XBP1* splicing or RIDD independently. While such strategies efficiently suppress *XBP1* splicing, they simultaneously affect additional IRE1-dependent outputs, including RIDD. In contrast, whether selective manipulation of the *XBP1* RNA substrate can uncouple *XBP1* splicing from other IRE1 outputs remains incompletely understood. We therefore hypothesized that targeting the *XBP1* splicing substrate could selectively interfere with unconventional *XBP1* splicing while preserving IRE1 activation, kinase activity and RIDD.

Here, we developed antisense morpholino oligonucleotides targeting the *XBP1* splice substrate and identified a morpholino that selectively suppresses *XBP1* splicing across multiple ER stress paradigms. Mechanistically, M2 reduced *XBP1s* production without detectable effects on IRE1 phosphorylation, recombinant IRE1 kinase activity or cleavage of a minimal fluorogenic IRE1 substrate, while altering the processing pattern of the native *XBP1* transcript. Together, our findings establish a substrate-directed strategy for modulation of the IRE1–XBP1 branch of the UPR and demonstrate that targeting the *XBP1* splice substrate is sufficient to redirect IRE1-mediated RNA processing without direct inhibition of IRE1.

## RESULTS

### Screening of antisense morpholinos targeting the *XBP1* splice substrate identifies a potent inhibitor of *XBP1* splicing

To identify an antisense morpholino capable of selectively suppressing IRE1-mediated *XBP1* splicing, we evaluated a panel of four splice-site-targeting morpholinos (M1-M4) and one translation-blocking morpholino (T1) designed against the XBP1 transcript (**Figure 1**). HeLa cells were transfected with individual morpholinos and subsequently exposed to tunicamycin (Tm) to induce ER stress and activate *XBP1* splicing. Tm treatment strongly increased *XBP1s* expression in control cells (**Figure 2A**). Among the splice-site-targeting morpholinos, M2 produced the most pronounced suppression of *XBP1s* mRNA, reducing XBP1s levels nearly to basal values, whereas M1, M3 and M4 exerted more modest inhibitory effects (**Figure 2A**). In contrast, the translation-blocking morpholino T1 did not reduce *XBP1s* transcript abundance and displayed XBP1s levels comparable to or higher than those observed in Tm-treated control cells (**Figure 2A**), consistent with its design to inhibit protein synthesis rather than RNA processing. Analysis of *XBP1u* expression revealed an opposite trend. Whereas Tm treatment substantially decreased the abundance of the unspliced transcript, reflecting active conversion of *XBP1u* into *XBP1s*, accumulation of *XBP1u* was observed in cells treated with splice-site-targeting morpholinos, particularly M2 (**Figure 2B**). This reciprocal decrease in *XBP1s* and increase in *XBP1u* is consistent with inhibition of IRE1-mediated splicing rather than transcript depletion. M2 treatment had a substantially greater effect on the distribution of *XBP1* splice variants than on total *XBP1* transcript abundance, suggesting that the primary consequence of M2 treatment was altered *XBP1* RNA processing rather than global transcript depletion (**Figure 2C**). In contrast, the translation-blocking morpholino T1 produced a distinct expression profile but failed to substantially reduce XBP1 protein abundance under the conditions tested (**Supplementary Figure S1**). Consistent with the reduction in *XBP1s* mRNA, M2 also significantly reduced stress-induced XBP1s protein accumulation (**Figure 2D**). Collectively, these data identify M2 as the most effective morpholino targeting the XBP1 splice substrate. The pronounced reduction in *XBP1s* abundance together with reciprocal accumulation of *XBP1u* is consistent with altered *XBP1* RNA processing rather than global depletion of *XBP1* transcripts. M2 was therefore selected for all subsequent mechanistic studies.

**Figure 1.**
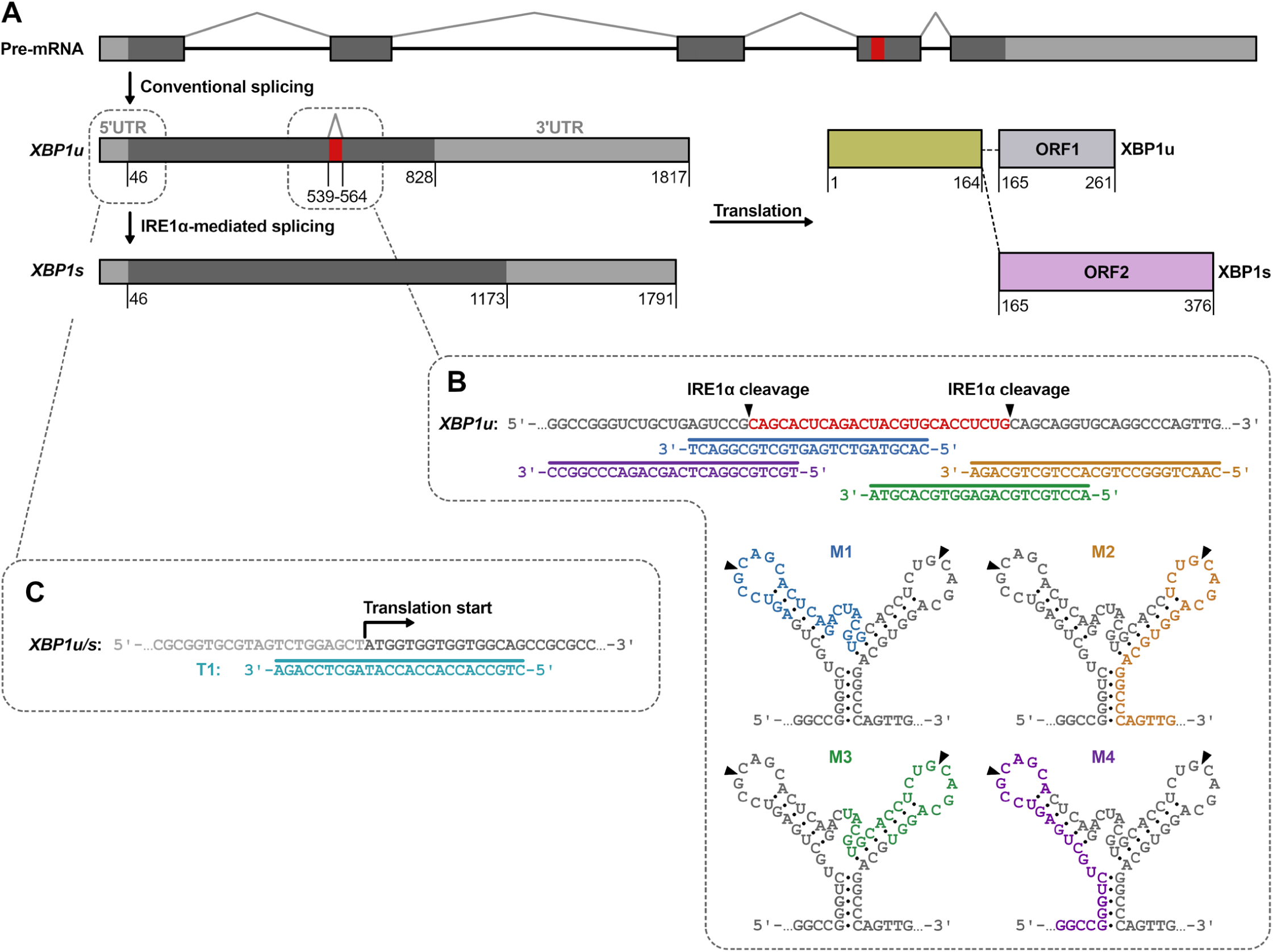
Design of antisense morpholino oligonucleotides targeting the *XBP1* splice substrate. **(A)** Schematic representation of conventional and unconventional splicing of *XBP1* mRNA. During ER stress, activated IRE1 excises a 26-nt intron from *XBP1u* mRNA, generating the spliced transcript *XBP1s*, which encodes the active transcription factor. The unconventional intron is indicated in red. **(B)** Design strategy of antisense morpholino oligonucleotides targeting the stem-loop structure encompassing the two canonical IRE1 cleavage sites within *XBP1u* mRNA. Four morpholinos (M1-M4) were designed to hybridize to distinct regions of the conserved *XBP1* splice substrate. The predicted binding sites of individual morpholinos relative to the IRE1 cleavage sites are indicated. Secondary structure predictions of the targeted region and the corresponding morpholino binding positions are shown. **(C)** Design of the translation-blocking morpholino T1 complementary to the region surrounding the *XBP1* translation initiation site. T1 was designed to inhibit XBP1 protein synthesis without directly targeting the *XBP1* splice region.

**Figure 2.**
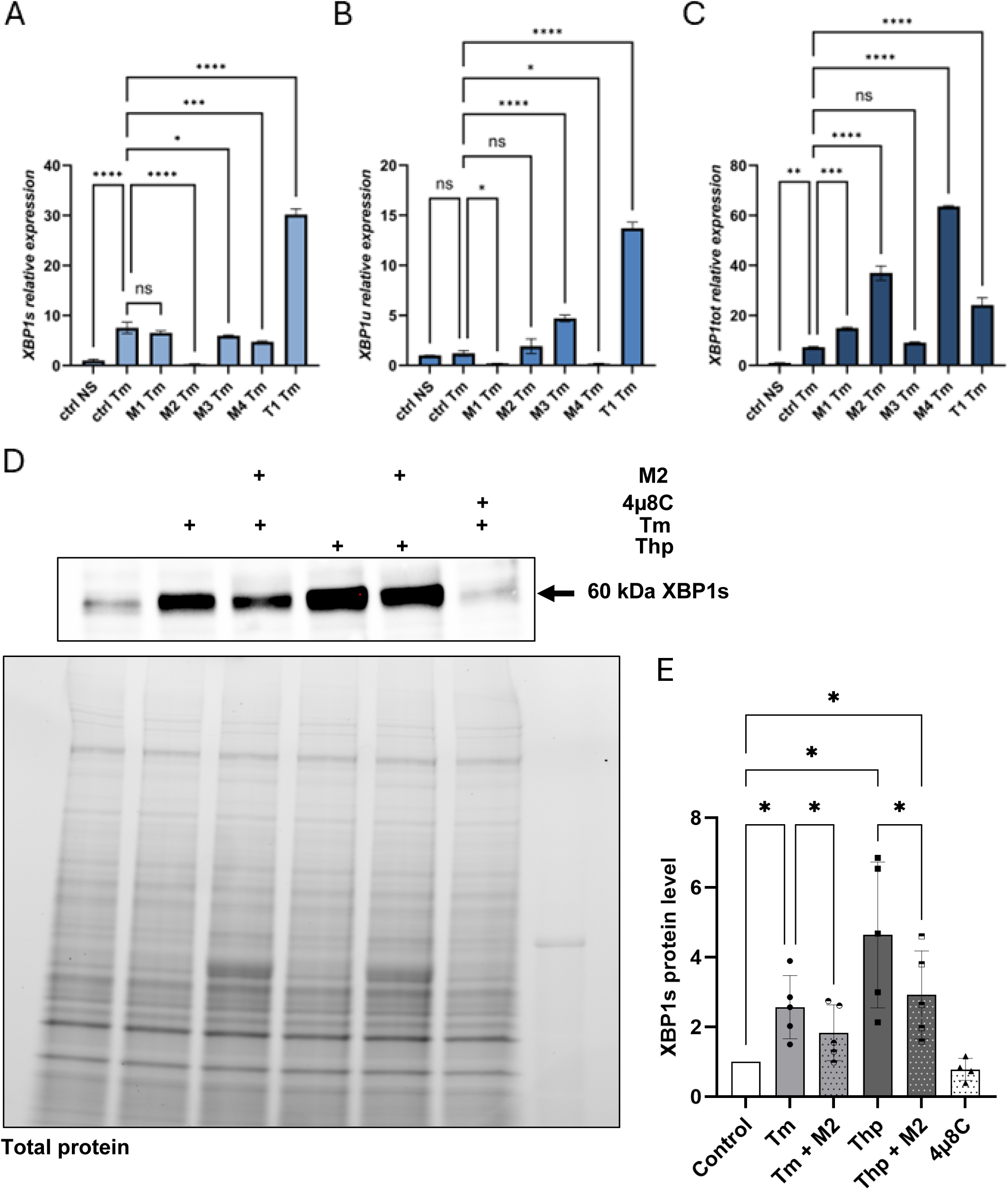
Identification of M2 as the most effective morpholino targeting the *XBP1* splice substrate. Relative expression of *XBP1s* (**A**), *XBP1u* (**B**) and total *XBP1* (**C**) mRNA in control and tunicamycin (Tm)-treated HeLa cells transfected with individual morpholino candidates (M1-M4 or T1), measured by RT-qPCR. Data are presented as mean ± SEM. Statistical significance was determined by one-way ANOVA with multiple-comparison testing: ns, not significant; P < 0.05 (), P < 0.01 (), P < 0.001 (), P < 0.0001 (****), n=3. (**D**) Analysis of XBP1s protein abundance following treatment with candidate morpholinos. Left panel: representative immunoblot showing XBP1s protein levels in HeLa cells exposed to tunicamycin (Tm) or thapsigargin (Thp) in the presence or absence of M2. Cells treated with the IRE1 RNase inhibitor 4μ8C served as a control for inhibition of XBP1s production. Total protein staining (TGX Stain-Free™ gel) is shown as a loading control. (**E**) Quantification of XBP1s protein levels relative to untreated control cells. Quantification was performed using five independent biological replicates (n=5) and data are presented as mean ± SEM (**Supplementary Figure S5**). Statistical significance was determined by one-way ANOVA followed by Fisher’s LSD multiple-comparison test; P < 0.05 (*).

### M2 suppresses stress-induced *XBP1* splicing in a dose-dependent manner

Following the identification of M2 as the most effective morpholino targeting the *XBP1* splice substrate, we further characterized its ability to suppress XBP1 splicing under ER stress conditions. Treatment of HeLa cells with tunicamycin (Tm) induced robust XBP1s expression, whereas transfection with M2 reduced *XBP1s* levels in a concentration-dependent manner (**Figure 3A**). Near-complete suppression of *XBP1s* accumulation was observed at 2 μM M2, while progressively weaker inhibition was detected at lower concentrations, confirming dose-dependent inhibition of *XBP1* splicing. To evaluate the persistence of this effect, we next examined *XBP1* splicing kinetics during ER stress. In control cells, *XBP1s* expression increased rapidly following Tm treatment and reached maximal levels approximately 6 h after stress induction (**Figure 3B**). In contrast, M2-treated cells exhibited markedly reduced *XBP1s* accumulation throughout the time course. Significant inhibition was observed at 6 h and 8 h and remained apparent at later time points, demonstrating sustained suppression of stress-induced XBP1s production. We next investigated whether reduced XBP1s generation was accompanied by accumulation of the unspliced transcript. Treatment with the IRE1 RNase inhibitor 4μ8C resulted in pronounced accumulation of *XBP1u* mRNA throughout the experiment, consistent with inhibition of XBP1 mRNA processing (**Figure 3C**). In contrast, M2 produced only modest changes in *XBP1u* abundance. Following prolonged ER stress, *XBP1u* levels in M2-treated cells remained similar to those observed in control cells and substantially lower than those detected following 4μ8C treatment. Thus, despite efficiently reducing *XBP1s* generation, M2 did not induce the marked accumulation of *XBP1u* typically associated with pharmacological inhibition of IRE1. Collectively, these findings demonstrate that M2 suppresses stress-induced *XBP1* splicing in a dose-dependent manner and maintains its inhibitory activity throughout the ER stress response. Unlike direct inhibition of IRE1 by 4μ8C, M2 reduces *XBP1s* accumulation while producing only a moderate increase in *XBP1u* abundance, indicating that the cellular consequences of M2 treatment differ from those associated with pharmacological inhibition of IRE1. These results establish M2 as a suitable tool for subsequent mechanistic studies of XBP1 RNA processing.

**Figure 3.**
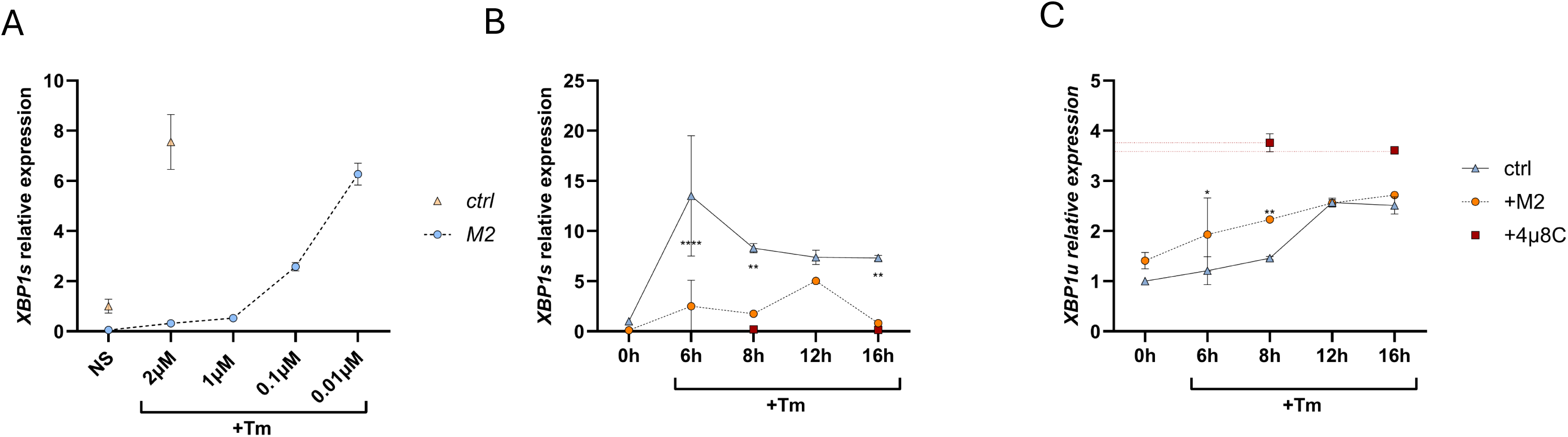
Functional validation of M2 as an inhibitor of *XBP1* mRNA splicing. (**A**) Dose-dependent inhibition of tunicamycin (Tm)-induced *XBP1s* expression by M2 in HeLa cells. (**B**) Time-course analysis of *XBP1s* expression following Tm treatment in control, M2-transfected and 4μ8C-treated HeLa cells. (**C**) Time-course analysis of *XBP1u* expression following Tm treatment in control, M2-transfected and 4μ8C-treated HeLa cells. Transcript levels were quantified by RT-qPCR and are presented as mean ± SEM. Statistical significance was determined by one-way ANOVA followed by Fisher’s LSD multiple-comparison test: ns, not significant; *P* < 0.05 (\**), P < 0.01 (**), P < 0.0001 (*\*\*\*), n=3.

### M2 selectively suppresses *XBP1* splicing under various ER stress conditions and cellular models

Having established M2 as an effective inhibitor of stress-induced *XBP1* splicing, we next examined whether its activity was restricted to a particular mode of ER stress induction. HeLa cells were exposed to a panel of mechanistically distinct ER stressors, including thapsigargin (Thp), dithiothreitol (DTT), brefeldin A (BFA) and the calpain inhibitor ALLN. As expected, each treatment induced substantial XBP1s expression relative to unstressed cells (**Figure 4A**). Under all conditions tested, transfection with M2 significantly reduced stress-induced *XBP1s* accumulation, indicating that the inhibitory effect of M2 is maintained independently of the ER stressor used. To further assess the robustness of this response, we evaluated the activity of M2 under conditions of pharmacological activation of the IRE1–XBP1 pathway. Treatment of HeLa cells with either tunicamycin or the selective IRE1 activator IXA4 markedly increased *XBP1s* expression compared with unstressed controls (**Figure 4B**). In both settings, *XBP1s* levels were significantly lower in M2-transfected cells than in the corresponding control groups. These results demonstrate that the inhibitory activity of M2 is retained under conditions of both ER stress-induced and pharmacological activation of *XBP1* splicing. We next investigated whether the activity of M2 could be reproduced in additional cellular models. In MDA-MB-231 cells, tunicamycin treatment induced sustained *XBP1s* expression throughout the analyzed time course (**Figure 4C**). Treatment with either M2 or 4μ8C was associated with reduced XBP1s accumulation at all examined time points. Similar results were obtained in MCF7 cells, where ER stress elicited a more pronounced induction of XBP1s expression (**Figure 4D**). In this model, M2 significantly reduced *XBP1s* levels throughout the time course, resulting in *XBP1s* expression levels comparable to those observed following 4μ8C treatment at later time points. Collectively, these findings demonstrate that M2 suppresses *XBP1s* accumulation across multiple ER stress paradigms and cellular models. The reproducibility of this effect in response to diverse ER stress stimuli, during pharmacological activation of the IRE1–XBP1 pathway, and in distinct cancer cell lines supports the use of M2 in subsequent mechanistic studies.

**Figure 4.**
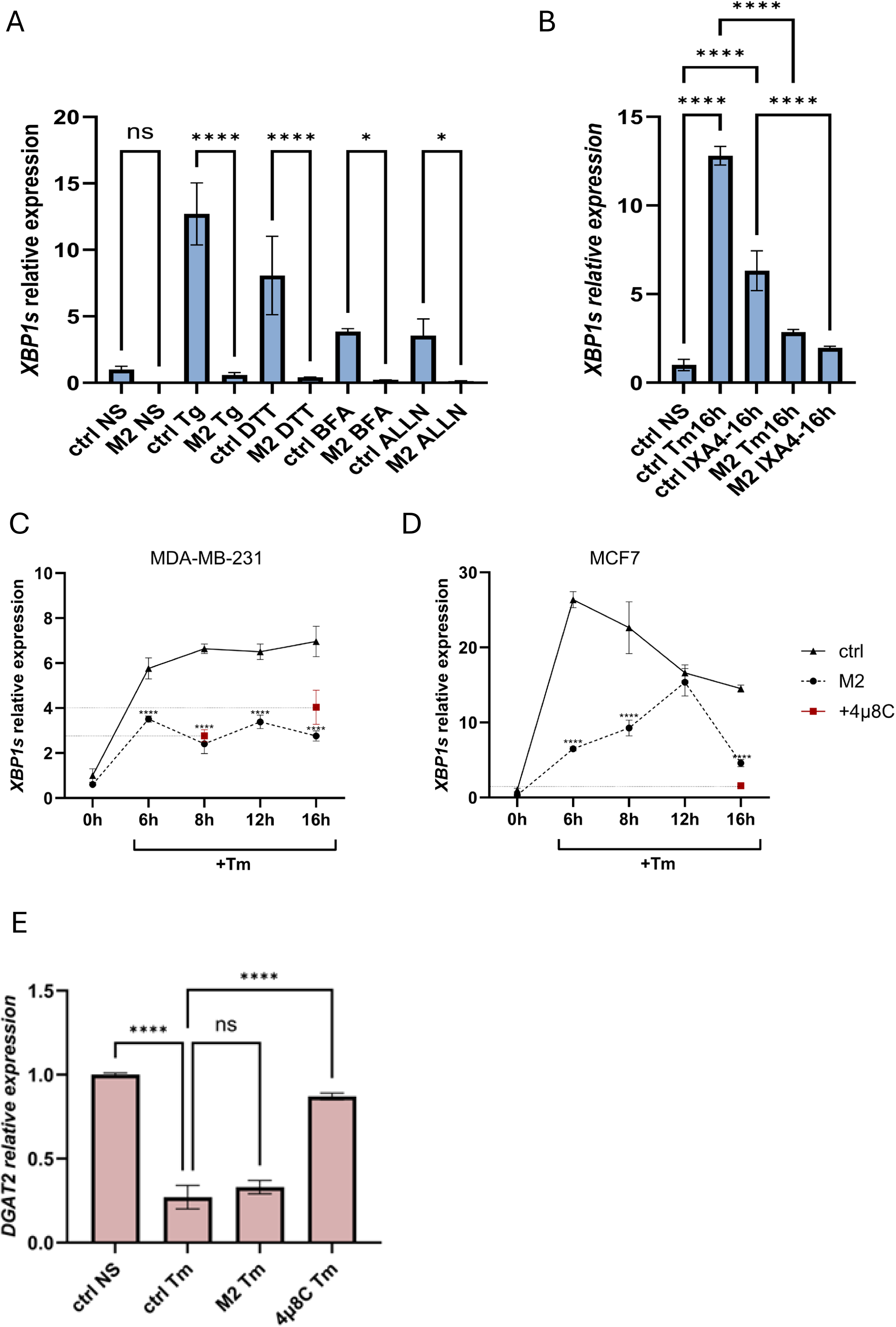
M2 suppresses *XBP1* splicing across multiple ER stress paradigms and cellular models. (**A**) Relative *XBP1s* expression in HeLa cells transfected with M2 and exposed to distinct ER stress inducers, including thapsigargin (Thp), dithiothreitol (DTT), brefeldin A (BFA) and the calpain inhibitor ALLN. (**B**) Relative *XBP1s* expression in control and M2-transfected HeLa cells following tunicamycin (Tm) treatment or pharmacological activation of IRE1 with IXA4. (**C**) Time-course analysis of *XBP1s* expression in MDA-MB-231 cells following tunicamycin treatment in the presence or absence of M2 or the IRE1 inhibitor 4μ8C. (**D**) Time-course analysis of *XBP1s* expression in MCF7 cells following tunicamycin treatment in the presence or absence of M2 or the IRE1 inhibitor 4μ8C. Transcript levels were determined by RT-qPCR and are presented as mean ± SEM. Statistical significance was determined by one-way ANOVA followed by Fisher’s LSD multiple-comparison test: ns, not significant; P < 0.05 (*), P < 0.01 (**), P < 0.001 (***), P < 0.0001 (****) n=3.(**E**) **M2 does not prevent stress-induced downregulation of the RIDD substrate *DGAT2.*** Relative *DGAT2* mRNA levels were measured by RT-qPCR in HeLa cells under basal conditions (ctrl NS) or following tunicamycin (Tm)-induced ER stress in the presence or absence of M2 or the IRE1 RNase inhibitor 4μ8C. Tm treatment resulted in a marked reduction in *DGAT2* transcript abundance. M2 did not significantly affect *DGAT2* expression compared with Tm-treated control cells, whereas 4μ8C substantially restored *DGAT2* levels. Data are presented as mean ± SEM. Statistical significance was assessed by one-way ANOVA followed by Fisher’s LSD multiple-comparison test: ns, not significant; P < 0.0001 (****) n=3.

### M2 does not alter stress-induced degradation of the RIDD substrate *DGAT2*

To determine whether M2-mediated suppression of *XBP1* splicing is accompanied by changes in RIDD, we examined the expression of *DGAT2* under ER stress [13]. Treatment of HeLa cells with Tm resulted in a significant reduction in *DGAT2* mRNA abundance relative to unstressed cells, consistent with stress-induced degradation of the transcript (**Figure 4E**).

Transfection with M2 did not significantly affect *DGAT2* expression in Tm-treated cells. *DGAT2* transcript levels remained similarly reduced in both control and M2-transfected cells, and no significant difference was detected between these groups (**Figure4E**). In contrast, treatment with the selective IRE1 RNase inhibitor 4μ8C largely prevented *DGAT2* degradation, resulting in transcript levels significantly higher than those observed in either control or M2-treated cells under ER stress conditions. Thus, although both M2 and 4μ8C reduce XBP1s production, they exhibit distinct effects on *DGAT2* expression. Whereas pharmacological inhibition of IRE1 with 4μ8C prevented the stress-induced reduction of *DGAT2* mRNA, M2 did not measurably alter *DGAT2* transcript abundance compared with Tm-treated control cells. These observations indicate that XBP1 splicing can be selectively suppressed without reproducing at least one well-established consequence of direct IRE1 inhibition and support substrate-directed modulation as a complementary strategy to enzyme-directed inhibition.

### M2 suppresses *XBP1* splicing without detectable effects on IRE1 activation or catalytic activity

The ability of M2 to suppress *XBP1* splicing raised the question of whether its activity results from direct inhibition of IRE1. To address this possibility, we first examined the phosphorylation status of IRE1 during ER stress. HeLa cells transfected with either control morpholino or M2 were exposed to Thp or Tm, and IRE1 phosphorylation was analyzed using Phos-tag SDS-PAGE and western blot. Both stressors induced the appearance of phosphorylated IRE1 species, whereas M2 treatment did not detectably alter the extent of IRE1 phosphorylation relative to control cells (**Figure 5A**).

**Figure 5.**
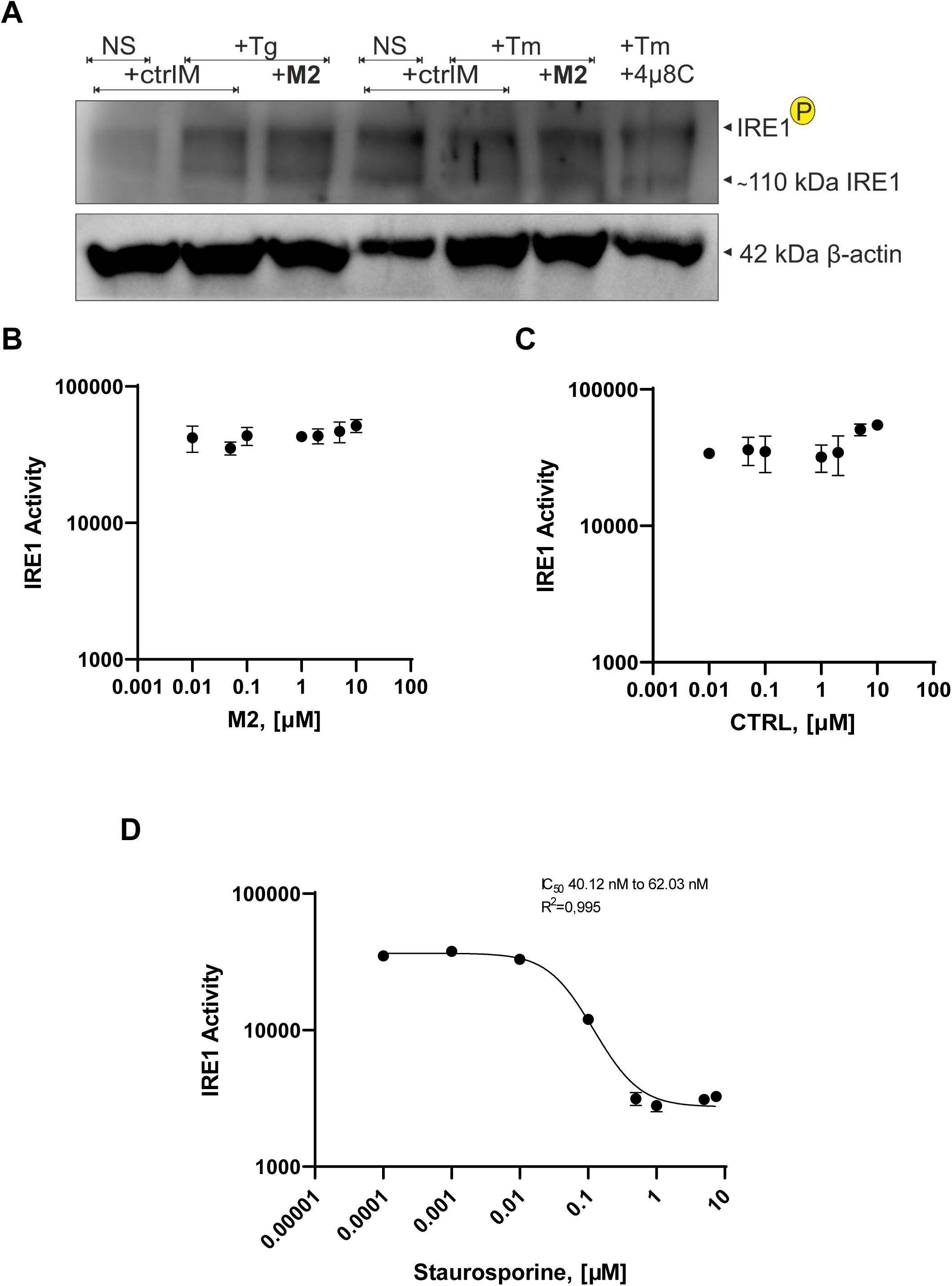
M2 suppresses *XBP1* splicing without affecting IRE1 activation or kinase activity. (**A**) Phos-tag SDS-PAGE followed by immunoblot analysis of total IRE1 and phosphorylated IRE1 (P-IRE1) in HeLa cells transfected with control morpholino (ctrlM) or M2 and subsequently exposed to thapsigargin (Thp) or tunicamycin (Tm). Cells treated with the IRE1 RNase inhibitor 4μ8C were included as a control. β-actin was used as a loading control. (**B**) Recombinant IRE1 kinase activity measured using the ADP-Glo kinase assay in the presence of increasing concentrations of M2. (**C**) Recombinant IRE1 kinase activity measured in the presence of a control morpholino (ctrlM). (**D**) Dose-response analysis of recombinant IRE1 kinase activity in the presence of staurosporine, which served as a positive control for kinase inhibition. Staurosporine inhibited IRE1 kinase activity in a concentration-dependent manner with an estimated IC₅₀ of 44.2 nM (95% CI 40.1–62.0 nM; R² = 0.995). Data are presented as mean ± SEM. Representative cropped blots are shown in Figure 5A; full-length immunoblots are provided in **Supplementary Figure S4**.

We next investigated whether M2 directly affects IRE1 kinase activity. Recombinant IRE1 was analyzed using an ADP-Glo kinase assay in the presence of increasing concentrations of M2. No reduction in kinase activity was observed across the concentration range tested for M2 (**Figure 5B**). Similarly, control morpholino and additional splice-site-targeting morpholinos (M1, M3 and M4) had no detectable effect on recombinant IRE1 kinase activity (**Figure 5C** and **Supplementary Figure S2**). As a positive control, the assay was performed in the presence of staurosporine, a broad-spectrum kinase inhibitor. Staurosporine inhibited recombinant IRE1 kinase activity in a concentration-dependent manner (IC₅₀ = 44.2 nM, 95% CI 40.1–62.0 nM; **Figure 5D**), confirming the sensitivity of the assay for detection of IRE1 kinase inhibition. Collectively, these findings indicate that M2 suppresses *XBP1* splicing without measurably affecting IRE1 phosphorylation or kinase activity. Together with the *DGAT2* data presented above, these results further distinguish the cellular consequences of M2 treatment from those associated with direct pharmacological inhibition of IRE1. Because M2 did not affect IRE1 phosphorylation or kinase activity, we next asked whether it might suppress *XBP1* splicing through direct inhibition of IRE1 RNase activity.

Next, to determine whether M2 suppresses XBP1 splicing through direct inhibition of IRE1 RNase activity, recombinant IRE1 was analyzed using a fluorogenic RNA cleavage assay based on a minimal XBP1-derived stem-loop substrate previously developed for selective measurement of IRE1 RNase activity [14, 15]. In this assay, cleavage of the fluorogenic RNA substrate results in increased fluorescence and provides a direct measure of IRE1-mediated RNase activity. Across the tested concentration range, M2 did not produce detectable inhibition of recombinant IRE1 RNase activity (**Figure 6A**). RNase activity remained comparable to control reactions even at the highest concentrations tested. As a positive control, reactions were performed in the presence of the selective IRE1 RNase inhibitor MKC8866. In contrast to M2, MKC8866 suppressed recombinant IRE1 RNase activity in a concentration-dependent manner (**Figure 6B**), confirming the sensitivity and dynamic range of the assay for detection of RNase inhibition. Importantly, the fluorogenic substrate used in this assay consists of a minimal IRE1-recognition stem-loop rather than the native XBP1 splice region targeted by M2. The absence of detectable inhibition therefore argues against a mechanism involving global suppression of IRE1 RNase activity. Collectively, these findings indicate that M2 does not function as a general inhibitor of IRE1 RNase activity. Rather, suppression of XBP1s production appears to result from altered processing of the native XBP1 substrate.

**Figure 6.**
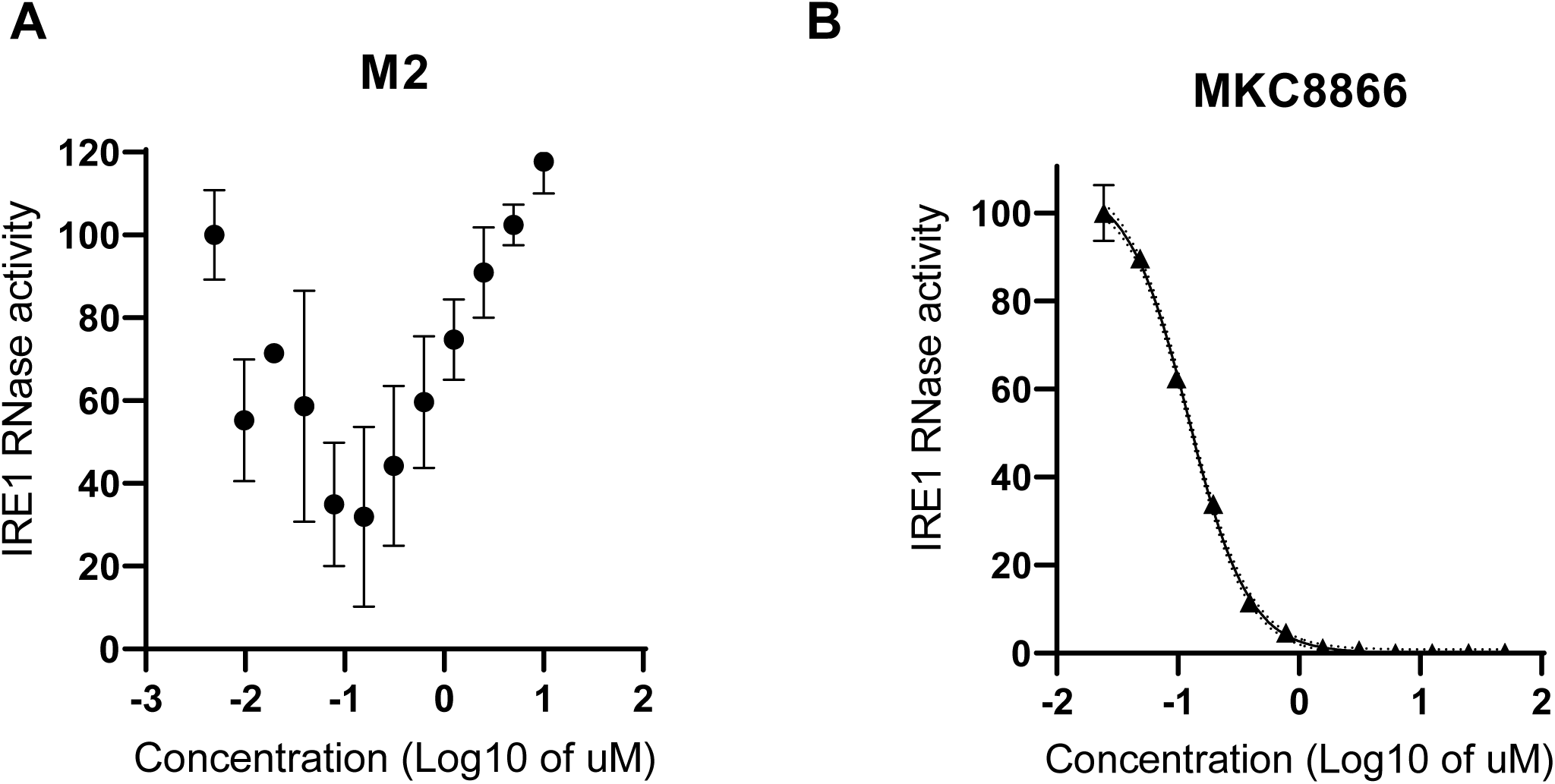
M2 does not inhibit cleavage of a minimal fluorogenic IRE1 RNA substrate. (**A**) Recombinant IRE1 RNase activity measured using a fluorogenic RNA cleavage assay containing a minimal *XBP1*-derived stem-loop substrate. Cleavage of the substrate separates the fluorophore from the quencher, resulting in increased fluorescence. M2 did not produce detectable inhibition of IRE1 RNase activity across the concentration range tested. (**B**) Dose-response analysis of recombinant IRE1 RNase activity in the presence of the selective IRE1 RNase inhibitor MKC8866, used as a positive control. MKC8866 inhibited substrate cleavage in a concentration-dependent manner, confirming the sensitivity and dynamic range of the assay for detection of IRE1 RNase inhibition. Data are presented as mean ± SEM, n=3.

### M2-mediated suppression of *XBP1* splicing is associated with increased persistence of *XBP1u*-associated RNA species

To determine whether the reduction in *XBP1s* abundance observed following M2 treatment could be attributed to altered transcript stability, we examined the decay kinetics of *XBP1u* and *XBP1s* mRNAs following transcriptional arrest with actinomycin D under ER stress conditions. Transcript abundance was quantified by RT-qPCR, normalized to levels at the start of the chase experiment, and fitted using an exponential decay model to estimate transcript half-lives. Under control conditions, RNA detected by the *XBP1u assay* exhibited an estimated half-life of approximately 4.0 h (**Figure 7A**). In contrast, *XBP1u* decay was markedly delayed in M2-treated cells, resulting in an estimated half-life of approximately 6.8 h. Thus, M2 treatment was associated with substantially prolonged persistence of RNA species containing the XBP1u assay target region. In contrast, only minor differences in *XBP1s* decay kinetics were observed. The estimated half-life of *XBP1s* increased from approximately 1.9 h in control cells to approximately 2.3 h following M2 treatment (**Figure 7B**). The overall decay profiles of *XBP1s* were largely comparable under both conditions, indicating that M2 had only a limited effect on the stability of the spliced transcript. As a control for transcript stability measurements, *GAPDH* mRNA turnover was analyzed in parallel. As expected, *GAPDH* displayed substantially slower decay kinetics, with an estimated half-life of approximately 7.9 h (**Figure 7C**), consistent with its established stability and previous report*s* [16, 17].

**Figure 7.**
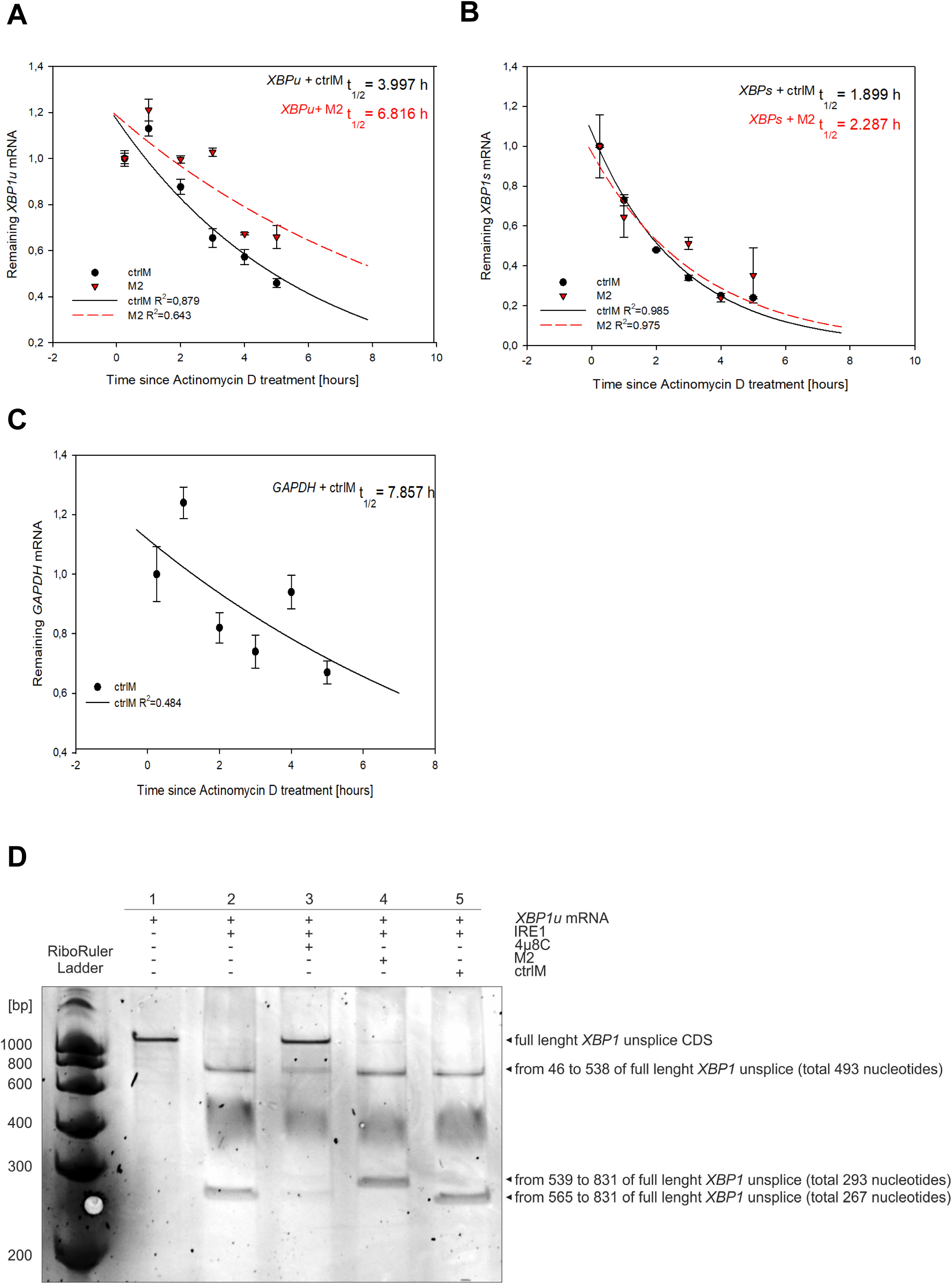
M2-mediated suppression of *XBP1* splicing is associated with increased persistence of *XBP1u*-associated RNA species. (**A**) Decay kinetics of RNA species detected by the *XBP1u* assay following transcriptional inhibition with actinomycin D during tunicamycin (Tm)-induced ER stress in HeLa cells transfected with control morpholino (ctrlM) or M2. Transcript abundance was determined by RT-qPCR, normalized to levels at time 0, and fitted using an exponential decay model to estimate apparent half-life. (**B**) Decay kinetics of *XBP1s* mRNA following transcriptional inhibition with actinomycin D during tunicamycin (Tm)-induced ER stress in HeLa CCL-2 cells transfected with ctrlM or M2. Transcript abundance was determined by RT-qPCR, normalized to levels at time 0, and fitted using an exponential decay model to estimate transcript half-life. **(C)** Decay kinetics of *GAPDH* mRNA analysed in parallel as a control transcript for RNA stability measurements. Data are presented as mean ± SEM. Solid and dashed lines represent exponential decay fitting used to calculate transcript half-lives, n=3 Estimated half-lives (t½) and goodness-of-fit values (R²) are indicated on the graphs. (**D**) **M2 alters the pattern of IRE1-mediated processing of XBP1 RNA *in vitro*.** An in vitro cleavage assay was performed using recombinant IRE1 and an XBP1u coding sequence RNA substrate. Cleavage products were separated by denaturing urea-PAGE and visualized with SYBR Gold. Lane 1, full-length XBP1u RNA substrate; lane 2, XBP1u RNA incubated with recombinant IRE1; lane 3, IRE1-mediated cleavage in the presence of the RNase inhibitor 4μ8C; lane 4, IRE1-mediated cleavage in the presence of M2; lane 5, IRE1-mediated cleavage in the presence of a control morpholino (ctrlM). Recombinant IRE1 generated the expected cleavage products of the XBP1 RNA substrate, whereas 4μ8C abolished RNA cleavage. M2 altered the cleavage pattern and promoted accumulation of a higher-molecular-weight RNA species not observed in control reactions.

Importantly, M2 did not accelerate the decay of either *XBP1* transcript. Instead, M2 markedly prolonged the persistence of *XBP1u* while exerting only a limited effect on *XBP1s* stability. Instead, M2 markedly increased the persistence of RNA detected by the *XBP1u* assay while exerting only a limited effect on *XBP1s* stability. These findings indicate that the reduction in *XBP1s* abundance observed following M2 treatment cannot be explained by enhanced transcript degradation. Rather, the data are consistent with altered processing of the *XBP1*splice substrate accompanied by increased persistence of RNA species containing the *XBP1u* region.

### M2 alters the pattern of IRE1-mediated *XBP1* RNA processing *in vitro*

To determine whether M2 directly affects processing of the *XBP1* substrate by IRE1, we performed an *in vitro* cleavage assay using recombinant IRE1 and an *XBP1u* RNA substrate. Cleavage products were separated by denaturing urea-PAGE and visualized by SYBR Gold staining (**Figure 7D**). As expected, recombinant IRE1 generated the characteristic cleavage products of the *XBP1* splice substrate, whereas the selective IRE1 RNase inhibitor 4μ8C completely abolished RNA cleavage (**Figure 7D**, **lanes 2** and **3**). Addition of a control morpholino (ctrlM) did not alter the cleavage pattern (**Figure 7D**, **lane 5**). In contrast, M2 altered the pattern of IRE1-mediated RNA processing and promoted accumulation of a higher-molecular-weight RNA species not observed in control reactions (**Figure 7D**, **lane 4**). Importantly, RNA cleavage was not abolished, as discrete cleavage products remained detectable. Thus, unlike 4μ8C, M2 did not phenocopy direct inhibition of IRE1 but instead modified the processing pattern of the *XBP1* substrate. Collectively, these findings demonstrate that M2 alters *XBP1* RNA processing by IRE1 *in vitro*. Together with the preserved IRE1 phosphorylation, kinase activity and fluorogenic RNase activity observed in other assays, these results support a model in which M2 suppresses XBP1s production through modulation of *XBP1* RNA processing rather than direct inhibition of IRE1.

## DISCUSSION

The IRE1–XBP1 branch of the unfolded protein response is most commonly manipulated through the modulation of IRE1 catalytic activity. While such an approach effectively suppress *XBP1* splicing, they simultaneously affect additional outputs of IRE1 signaling, including RIDD [10–12]. In the present study, we explored an alternative strategy based on direct targeting of the *XBP1* splicing substrate. Using antisense morpholino oligonucleotides directed against the *XBP1* splicing region, we identified M2 as a potent suppressor of XBP1s production and demonstrate that *XBP1* splicing can be selectively inhibited without detectable effects on IRE1 phosphorylation, kinase activity, cleavage of a minimal fluorogenic IRE1 substrate, or stress-induced downregulation of the RIDD substrate *DGAT2*.

The inhibitory activity of M2 proved robust across multiple experimental settings. M2 reduced *XBP1s* accumulation in response to mechanistically distinct ER stressors, including tunicamycin, thapsigargin, dithiothreitol, brefeldin A and ALLN, and remained effective following pharmacological activation of the IRE1–XBP1 pathway with IXA4. Furthermore, comparable effects were observed in multiple cellular backgrounds, including HeLa, MDA-MB-231 and MCF7 cells. These findings indicate that the activity of M2 is not restricted to a particular mode of ER stress induction or cellular context and support the general applicability of the observed phenotype. A central observation of this study is that suppression of *XBP1* splicing by M2 is not accompanied by several molecular consequences typically associated with direct inhibition of IRE1. In contrast to the selective IRE1 RNase inhibitor 4μ8C, M2 neither altered stress-induced IRE1 phosphorylation nor affected recombinant IRE1 kinase activity. Moreover, M2 did not prevent stress-induced downregulation of *DGAT2,* a well-characterized RIDD substrate, whereas 4μ8C restored *DGAT2* expression. Although analysis of a single RIDD target does not permit a general conclusion regarding the entirety of RIDD activity, these findings indicate that the cellular consequences of M2 treatment differ from those observed following pharmacological inhibition of IRE1. Thus, our data indicate that *XBP1* splicing can be selectively modulated without recapitulating the molecular consequences associated with inhibition of IRE1 catalytic activity.

One notable consequence of M2 treatment was the increased persistence of RNA species detected by the *XBP1u* assay. M2 increased the apparent half-life of these transcripts from approximately 4.0 h to 6.8 h, whereas only minor changes were observed for *XBP1s*. These data indicate that inhibition of XBP1s production occurs despite prolonged persistence of RNA containing the *XBP1u* region and therefore cannot be explained by enhanced RNA degradation.

The most direct mechanistic insight was provided by the in vitro cleavage assay. Recombinant IRE1 generated the expected cleavage products from *XBP1u* RNA, whereas 4μ8C completely abolished cleavage. In contrast, M2 produced a distinct processing pattern characterized by the accumulation of a higher-molecular-weight RNA species indicative of altered processing of the *XBP1* splice substrate. Importantly, cleavage activity was not eliminated. Instead, the data suggest that M2 interferes with productive processing of the *XBP1* substrate, resulting in accumulation of a processing intermediate while preserving detectable IRE1-mediated cleavage activity. These observations are therefore more consistent with altered substrate processing than with direct inhibition of the enzyme itself.

Importantly, M2 failed to inhibit cleavage of a minimal fluorogenic *XBP1*-derived substrate used for direct measurement of IRE1 RNase activity (**Figure 6**), whereas the selective IRE1 inhibitor MKC8866 produced robust suppression of substrate cleavage. Because this assay employs a minimal synthetic XBP1 stem-loop substrate [14, 15] rather than the extended native *XBP1* splice region targeted by M2, the observed lack of inhibition provides additional evidence against a mechanism involving direct suppression of IRE1 RNase activity. Thus, M2 alters processing of the native *XBP1* substrate without detectably affecting cleavage of a minimal IRE1-recognition substrate, further supporting a substrate-directed rather than enzyme-directed mechanism of action. Together, the preservation of IRE1 phosphorylation, kinase activity and fluorogenic RNase activity, combined with altered processing of the native *XBP1* substrate, supports a model in which M2 acts through substrate-directed modulation of *XBP1* RNA processing rather than direct inhibition of IRE1.

An additional observation supporting a substrate-directed mechanism was the marked functional heterogeneity among the splice-site-targeting morpholinos. Despite being designed to hybridize within the same general *XBP1* stem-loop region, M1-M4 differed substantially in their ability to suppress *XBP1s* production (**Figure 2D** and **Supplementary Figure S3**). In particular, M2 consistently exhibited greater inhibitory activity than the remaining morpholinos under both Tm-and Thp-induced ER stress conditions. These findings indicate that productive XBP1 processing is highly sensitive to the precise location of substrate perturbation and further support the notion that local structural features of the XBP1 splice substrate contribute to efficient IRE1-mediated processing.

Our findings are compatible with previous studies demonstrating that efficient *XBP1* splicing depends on highly specialized interactions between IRE1 and its RNA substrate. Earlier work showed that localization, structural organization and productive engagement of XBP1u are essential determinants of unconventional splicing [4, 5]. In parallel, Tam and colleagues demonstrated that *XBP1* splicing and RIDD possess distinct substrate requirements and catalytic features, indicating that individual outputs of IRE1 signaling can be mechanistically separated [12]. Our data extend this concept by showing that modulation of the substrate itself is sufficient to suppress *XBP1s* production without reproducing key molecular effects associated with direct inhibition of IRE1. Notably, the strategy described here differs conceptually from conventional approaches targeting IRE1. Rather than suppressing the catalytic activity of the enzyme, M2 acts at the level of the RNA substrate undergoing unconventional splicing. This distinction may prove useful for dissecting biological processes that depend specifically on XBP1s while minimizing perturbation of other measured outputs associated with activated IRE1. In this context, substrate-directed modulation and enzyme-directed inhibition should be viewed as complementary experimental approaches rather than interchangeable strategies [12, 18].

Several limitations should be acknowledged. First, the interaction between M2 and the *XBP1* splice region has not been mapped at nucleotide resolution. Second, the identity of the intermediate cleavage product observed in vitro was inferred from fragment size rather than directly confirmed by sequencing. Finally, although *DGAT2* analysis argues against disruption of at least one established RIDD event, analysis of additional endogenous RIDD substrates will be required to define the broader impact of M2 on IRE1 RNase outputs. Future studies combining structural approaches, RNA footprinting and high-resolution mapping of cleavage products should provide a more detailed understanding of how morpholino binding alters processing of the XBP1 splice substrate.

In summary, we describe a substrate-directed strategy for suppression of *XBP1* splicing. Unlike conventional IRE1 inhibitors, M2 reduces XBP1s production without detectable effects on IRE1 phosphorylation, IRE1 kinase activity, cleavage of a minimal fluorogenic IRE1 RNA substrate, or stress-induced downregulation of the RIDD substrate *DGAT2*. Mechanistically, M2 alters the pattern of IRE1-mediated processing of *XBP1* RNA and is associated with increased persistence of RNA species detected by the *XBP1u* assay. Together, these findings demonstrate that selective targeting of the *XBP1* splice substrate is sufficient to suppress *XBP1s* production through substrate-directed modulation of *XBP1* RNA processing rather than direct inhibition of IRE1 and provide a framework for investigating the biological functions of the IRE1–XBP1 pathway independently of IRE1 catalytic inhibition.

## METHODS

### Morpholino design

We designed antisense morpholino oligonucleotides (M1-M4) to target the highly conserved stem-loop structure within the *XBP1u* mRNA that contains the two canonical IRE1α cleavage sites required for unconventional splicing. Candidate sequences were selected to hybridize with distinct regions surrounding the cleavage motifs and the predicted stem-loop structure, thereby allowing experimental perturbation of the XBP1 splice substrate. T1 morpholino was designed to target the XBP1 translation initiation region encompassing the AUG start codon. A morpholino oligonucleotide targeting a human β-globin intron mutation associated with β-thalassemia was used as a negative control (ctrlM). The sequences of all morpholino oligonucleotides used in this study are listed in Table 1. All antisense morpholinos were synthesized and purchased from Gene Tools, LLC (Philomath, OR, USA).

**Table 1.**
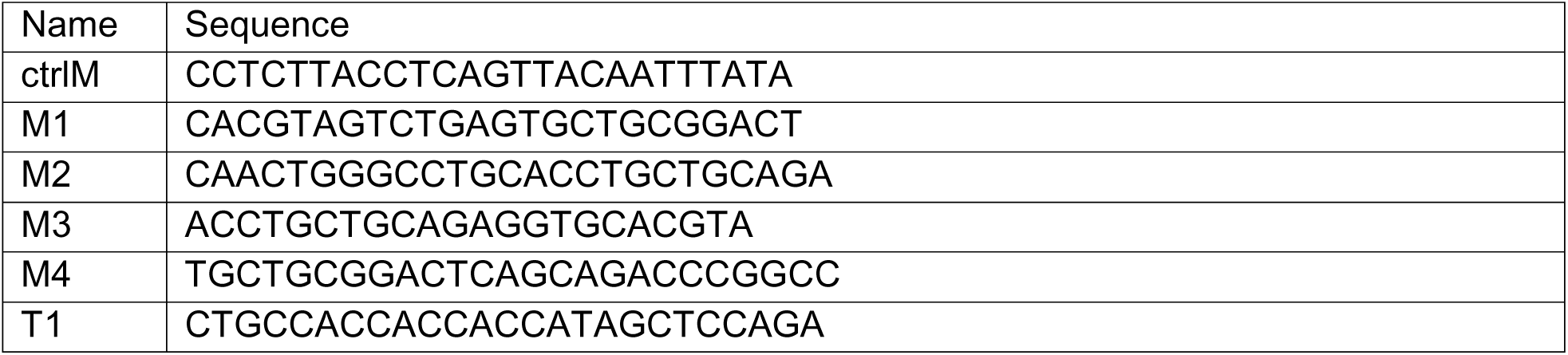
Morpholino oligonucleotides targeting the IRE1α cleavage region and XBP1 mRNA translation start site.

### Cell lines

The human cervical carcinoma HeLa (CCL-2™), the human breast adenocarcinoma MDA-MB-231 (HTB-26™), and the human breast adenocarcinoma MCF7 (HTB-22™) cell lines were obtained from the ATCC. The cells were maintained in Minimum Essential Modified Eagle’s Medium (MEM; Biowest, L0416) supplemented with 2 mM L-glutamine, 10% fetal bovine serum (FBS; Sigma-Aldrich, F7524) and antibiotics (100 μg/ml penicillin and 100 μg/ml streptomycin; Biowest, L0018). All cell lines were cultured at 37 °C in a humidified atmosphere containing 5% CO₂ and seeded at a density of 500,000 or 250,000 cells per well in 6-or 12-well plates one day prior to experimental procedures.

| **Cell culture** |  |  |
|---|---|---|
| fetal bovine serum, FBS | Sigma-Aldrich | F7524 |
| Minimum Essential Modified Eagle’s | Biowest | L0416 |
| penicillin and streptomycin solution | Biowest | L0018 |

### Morpholino Transfection

Cells were transfected with ctrlM, M1-M4 or T1 at the indicated final concentrations using Endo-Porter PEG (Gene Tools, LLC, Philomath, OR, USA) according to the manufacturer’s instructions (6 μl reagent per ml of culture medium). Morpholino delivery and ER stress induction were initiated simultaneously and maintained throughout the duration of the experiment. Cells were harvested at the indicated time points for subsequent RNA and protein analyses.

### Pharmacological induction of ER stress

Pharmacological induction of ER stress and activation of the unfolded protein response (UPR) were performed as previously described (12). Briefly, cells were treated with tunicamycin (Tm; 2.5 μg/ml; Sigma-Aldrich, T7765), thapsigargin (Thp; 50 nM; Sigma-Aldrich, T9033), dithiothreitol (DTT; 4 mM; Sigma-Aldrich, 43815), brefeldin A (BFA; 0.4 μg/ml; Sigma-Aldrich, B6542), or the calpain inhibitor ALLN (10 μM; Sigma-Aldrich, 208719) for the indicated time periods. Control cells received an equivalent volume of vehicle (DMSO; Sigma-Aldrich, D2650). To inhibit IRE1 RNase activity, cells were treated with 4μ8C (10 μM; Sigma-Aldrich, SML0949) or MKC8866 (indicated concentrations), both dissolved in DMSO. To activate the IRE1–XBP1 pathway independently of ER stress induction, cells were treated with IXA4 (10 μM; MedChemExpress, HY-139214) for 16 h

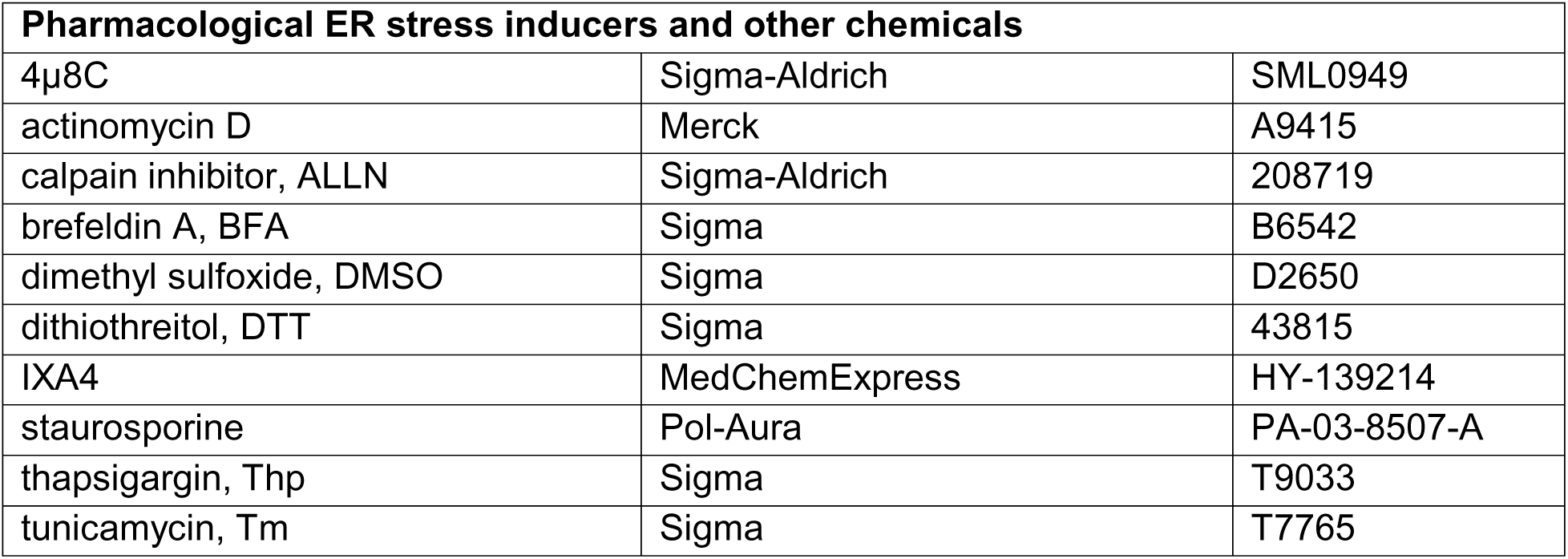

### Isolation of cellular RNA

Total RNA was extracted using the RNeasy Mini Kit (Qiagen, 74104) in accordance with the manufacturer’s instructions. Residual genomic DNA was removed from all samples, as recommended by the supplier. The concentration of the RNA was determined spectrophotometrically by measuring the absorbance at 260 nm. The purified RNA samples were stored at −20 °C until further analysis.

### Quantitative Real-Time PCR

Gene expression analysis was performed using the TaqMan RNA-to-CT™ 1-Step Kit (Applied Biosystems™, catalogue number 4392938), following the manufacturer’s instructions and the protocol previously described in reference[19]. This included a reverse transcription (RT) step carried out at 50 °C for 20 minutes. Relative transcript abundance was determined using the 2^−ΔΔCt^ method[20]. TATA-box binding protein (*TBP*; Hs00427620_m1) was used as the endogenous reference gene. For the mRNA stability analyzes eukaryotic 18S rRNA (*18S*; Hs03003631_g1) and glyceraldehyde-3-phosphate dehydrogenase (*GAPDH*; Hs03929097_g1) was used. All measurements were performed in technical triplicate, and expression levels were quantified from the resulting Ct values. The TaqMan probe assays used in this study included *DGAT2* (Hs01015913_m1), *XBP1tot* (Hs00231936_m1), *XBP1s* (Hs03929085_g1), *XBP1u* (Hs02856596_m1). The *XBP1u* assay targets a region upstream of the unconventional IRE1 splice sites and was therefore used to monitor RNA species containing the XBP1u assay target region. All probes were obtained from Life Technologies (Warsaw, Poland).

| **Gene expression assay** |  |  |
|---|---|---|
| *18S* | Thermo Fisher | Hs03003631_g1 |
| *DGAT2* |  | Hs01015913_m1 |
| *GAPDH* |  | Hs03929097_g1 |
| *TBP* |  | Hs00427620_m1 |
| *XBP1tot* |  | Hs00231936_m1 |
| *XBP1s* |  | Hs03929085_g1 |
| *XBP1u* |  | Hs02856596_m1 |

### Measurement of RNA half-life

The half-lives of *XBP1s*, *XBP1u*, and *GAPDH* transcripts were determined as previously described with minor modifications [21, 22]. 500,000 HeLa cells were seeded in 35-mm dishes and transfected with either ctrlM or M2. To induce ER stress, cells were treated with tunicamycin (Tm; 2.5 μg/mL) for 3 h prior to initiation of the decay assay. At the start of the time course, cells were simultaneously exposed to the IRE1 RNase inhibitor 4μ8C (10 μM) and the transcriptional inhibitor actinomycin D (5 μg/mL; Merck, A9415), which were maintained throughout the experiment. Total RNA was isolated at the indicated time points using the RNeasy Mini Kit (Qiagen, 7410). Transcript levels were quantified by TaqMan-based real-time PCR and normalized to endogenous *18S* RNA measured using a standard primer set. For each time point, data were obtained from three biological replicates across at least two independent experiments. Relative RNA abundance was calculated as the fold change relative to the initial time point (t = 0). RNA decay rates were estimated by exponential regression according to the equation C/C₀ = e⁻, where C and C₀ denote transcript abundance at time points t and t₀, respectively, and kd represents the decay constant, as previously described [22].

### Western Blots

The cells were lysed on ice for 15 minutes in RIPA buffer (Sigma-Aldrich, R0278), which was supplemented with a Protease Inhibitor cOmplete Mini (11836170001; Roche). The lysates were then homogenized by repeatedly passing them through a 27-gauge needle (20 times) and the insoluble material was removed by centrifugation at 15,000×g for 15 minutes at 4 °C. Protein concentrations were determined using the DC Protein Assay (Bio-Rad, 5000111), using bovine serum albumin (BSA; Biowest, P6155-KG) as a standard.

Equal amounts of protein were combined with 4× Laemmli sample buffer, heated at 95 °C for 5 min and separated using Criterion™ TGX Stain-Free™ 4–15% gradient gels (Bio-Rad, 5678084) or Mini-PROTEAN® TGX Stain-Free™ gels (Bio-Rad, 4568084). The proteins were then transferred onto PVDF membranes at 300 mA for 90 minutes at 4 °C. The membranes were then blocked overnight in TBS (20 mM Tris-HCl, 150 mM NaCl, pH 7.6) containing 5% BSA and 0.5% Tween-20 (IBI Scientific, IB07200). The primary antibodies were diluted in TBST (TBS supplemented with 3% BSA) and used at the following concentrations: anti-XBP1s (Cell Signaling Technology, #12782, 1:1000). After washing with TBST, the membranes were incubated with a horseradish peroxidase-conjugated goat anti-rabbit IgG secondary antibody (Bio-Rad, 1706515). Immunoreactive bands were visualized using SuperSignal™ West Femto substrate (Thermo Fisher Scientific, 34096) and chemiluminescent signals were acquired using a ChemiDoc™ XRS imaging system (Bio-Rad). Band intensities were quantified using Image Lab software v4.1 (Bio-Rad).

For the Phos-tag gel electrophoresis and λ-phosphatase treatment HeLa cells were harvested after 6 hours of treatment with Tm or under NS conditions, in 1× NEBuffer for Protein MetalloPhosphatases (PMP; containing 50 mM HEPES, 100 mM NaCl, 2 mM DTT and 0.01% Brij-35, with a pH of 7.5; New England Biolabs, P0753S), supplemented with 1 mM MnCl₂. Soluble protein fractions were treated either with λ-phosphatase (+ λ) or left untreated (− λ), according to the manufacturer’s instructions. Samples were prepared for Western blot analysis as described above, except that the proteins were mixed with 2× Laemmli buffer prior to being heated at 95 °C for 5 minutes. Protein separation was performed using SuperSep™ Phos-tag™ gels (FUJIFILM, 192-17401). Following transfer to PVDF membranes (GE Healthcare, 10600023) and an overnight block in 5% BSA in TBST, the membranes were incubated with anti-IRE1 (Abcam, ab37073), diluted 1:1000 in TBST containing 3% BSA. Signal detection and image acquisition were performed as described above.

| **Proteins isolation, handling and Western blot utilities** |  |  |
|---|---|---|
| λ phosphatase | Biolabs | P0753S |
| 7.5% TGX<sup>TM</sup> FastCast<sup>TM</sup> polyacrylamide gel | Bio-Rad | 161-0171 |
| 4× Laemmli buffer | Bio-Rad | 1610747 |
| anti-IRE1 | Abcam | ab37073 |
| anti-rabbit IgG secondary antibody | Bio-Rad | 1706515 |
| anti-XBP1s | Cell Signaling Technology | #12782 |
| Criterion TGX Stain-Free 4 – 15% gradient gels | Bio-Rad | 5678084 |
| DC Protein Assay | Bio-Rad | 5000111 |
| Dulbecco’s Phosphate Buffered Saline, PBS | Biowest | L0616 |
| Mini-PROTEAN TGX Stain-Free gels | Bio-Rad | 4568084 |
| Protease Inhibitor cOmplete Mini | Roche | 11836170001 |
| PVDF membrane | GE Healthcare | 10600023 |
| SuperSep Phos-tag gels | FujiFilm | 192-17401 |
| SuperSignal West Femto substrate | Thermo Fisher Scientific | 34096 |
| Tween-20 | IBI Scientific | IB07200 |

### *In vitro* IRE1 kinase assay

The effect of antisense morpholino oligonucleotides on IRE1 kinase activity was evaluated using the ADP-Glo™ Kinase Assay (Promega, V6930) with recombinant human IRE1 kinase (Promega, VA7144). Kinase reactions were performed in 1× Reaction Buffer A (40 mM Tris-HCl [pH 7.5], 20 mM MgCl₂, and 0.1 mg/mL BSA), supplemented with 50 μM DTT. Prior to inhibitor screening, IRE1 enzyme titration was performed to determine the amount of kinase yielding approximately 10% ATP-to-ADP conversion within the linear range of the assay. Based on these results, 8.2 ng of recombinant ERN1/IRE1α was used per reaction in all subsequent assays. The reaction mixture contained the following: recombinant IRE1, myelin basic protein (MBP; 0.1 μg/μL) as the substrate, ATP at a final concentration of 10 μM. Kinase reactions were assembled in white 96-well plates at a final volume of 25 μL and incubated for 60 minutes at RT. Morpholino oligonucleotides (ctrlM, M1-M4) were tested at final concentrations ranging from 0.01 to 10 μM. Staurosporine (Pol-Aura, PA-03-8507-A) was included as a positive control inhibitor and tested at concentrations ranging from 0.01 to 20 μM. The kinase reaction was initiated by the addition of IRE1 and performed in triplicate. After the kinase incubation, 25 μL of ADP-Glo™ Reagent was added to each well, and the plates were incubated for 40 minutes at RT to terminate the reaction and deplete the unconsumed ATP. Then, 50 μL of Kinase Detection Reagent was added, and the samples were incubated for an additional 30 minutes at RT. Luminescence was measured using a GloMax® Discover Microplate Reader (Promega) with a one-second integration time. Kinase activity was expressed as relative luminescence units (RLU), which correlate with ADP production and ATP consumption. Dose-response curves were generated by plotting IRE1 kinase activity against inhibitor concentration.

| **Luminescence assays** |  |  |
|---|---|---|
| ADP-Glo™ Kinase Assay | Promega | V6930 |
| IRE1 kinase | Promega | VA7144 |

### Minimal fluorogenic in vitro IRE1 RNase assay

Recombinant human IRE1α RNase activity was measured using a fluorogenic RNA cleavage assay based on a minimal XBP1-derived stem-loop substrate previously developed and validated for selective detection of IRE1-mediated RNA cleavage [15, 23]. The synthetic RNA substrate contained a fluorophore (Cy5, emission at 590 nm) and quencher positioned such that IRE1-dependent cleavage resulted in increased fluorescence. Recombinant IRE1α protein (0.6 µg/reaction) was incubated with the fluorogenic RNA substrate (1 µg) in reaction buffer (20 mM HEPES, pH 7.5; 1 mM MgOAc; 50 mM KOAc, supplemented with 20 mM ATP and 2 mM DTT; final DMSO ≤ 1%) according to the protocol [15, 23]. Fluorescence was monitored every minute for 25 min at 37 °C using a microplate reader (Tecan Infinite 200) and used as a direct measure of IRE1 RNase activity. To assess potential effects of morpholinos on IRE1 RNase function, recombinant IRE1α was incubated for 10 min at room temperature with increasing concentrations of M2 prior to addition of the fluorogenic substrate. The selective IRE1 RNase inhibitor MKC8866 was included as a positive control. Fluorescence values were normalized to vehicle-treated controls and expressed as relative IRE1 RNase activity. Because this assay utilizes a minimal synthetic *XBP1* stem-loop substrate rather than the extended native *XBP1* splice region targeted by the morpholino oligonucleotides, it was used to evaluate direct effects on IRE1 RNase activity independently of interactions with the full-length XBP1 splice substrate.

### Template preparation

To generate the *XBP1u* mRNA transcription template, a plasmid was constructed that contained the *XBP1u* coding sequence (CDS; nucleotides 49-834), preceded by the T7 Φ2.5 RNA polymerase promoter at the 5′ end. The EX-F0758-M02 vector (GeneCopoeia, Rockville, MD, USA) served as the cloning template. PCR amplification was performed using 20 ng of template DNA, 2X Taq Master Mix (New England Biolabs, M0270), and 0.5 μM of each forward and reverse primer (Table 2). The PCR conditions are summarized in Table 3.

**Table 2.**
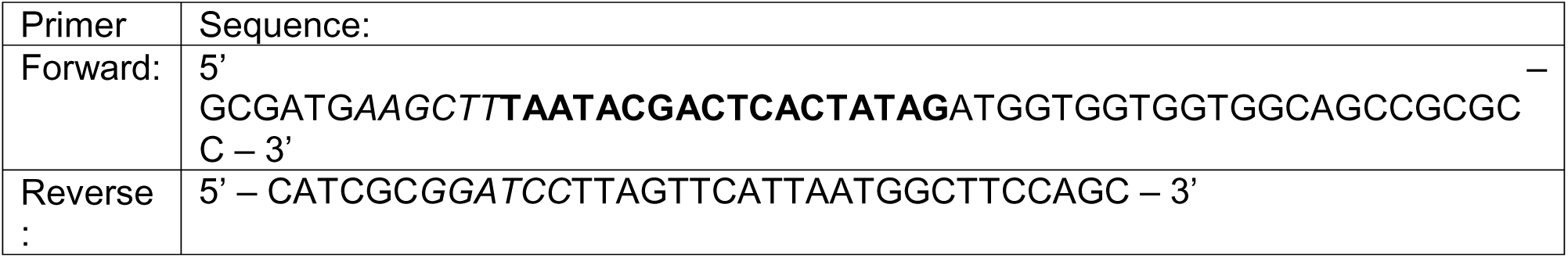
Primers used in the PCR reaction to amplify CDS of XBP1u. The promoter sequence for polymerase T7 Φ2.5 is in bold, and the restriction site for HindIII in the forward primer and for BamHI in the reverse primer is in italics. These sites enabled cloning into the pUC19 plasmid vector. All primers were synthesized by Genomed, Warsaw, Poland.

**Table 3.** PCR reaction programs for XBP1u CDS amplification.

|  | **CDS of *XBP1u*** | **Step** | **No. of cycles** |
|---|---|---|---|
| temperature [°C] | 95 | initial denaturation | - |
| time [s] | 120 |  |  |
| temperature [°C] | 95 | denaturation | X30 |
| time [s] | 25 |  |  |
| temperature [°C] | 68 | annealing |  |
| time [s] | 30 |  |  |
| temperature [°C] | 72 | extension |  |
| time [s] | 75 |  |  |
| time [s] | 45 |  |  |
| temperature [°C] | 72 | final extension | - |
| time [s] | 300 |  |  |
| temperature [°C] | 4 | hold |  |
| time [s] | ∞ |  |  |

The PCR products were purified using NucleoSpin Gel and PCR Clean-up columns (Macherey-Nagel, 740609.50S). Then, they were digested with *BamHI* (Thermo Scientific, FD0054) and *HindIII* (Thermo Scientific, FD0505), according to the manufacturers’ recommendations. Finally, the products were purified again prior to cloning. The resulting inserts were ligated into the *BamHI/HindIII*-linearized pUC19 vector at a 1.5:1 insert-to-vector molar ratio using T4 DNA ligase (Thermo Scientific, EL0014), following the supplier’s protocol. Competent *Escherichia coli* DH5α cells were transformed as previously described and plated on LB agar supplemented with 100 μg/mL ampicillin (BioShop Canada Inc., AMP201.100)[24]. Plates were incubated overnight at 37 °C. Single colonies were used to establish bacterial overnight cultures in 3 mL LB medium containing 100 μg/mL ampicillin at 37°C with shaking at 182 rpm. Plasmid DNA was subsequently isolated using the GeneMATRIX Basic DNA Purification Kit (EURx, E3545). The correct insertion and sequence integrity of the cloned construct were confirmed by sequencing performed by Genomed (Warsaw, Poland).

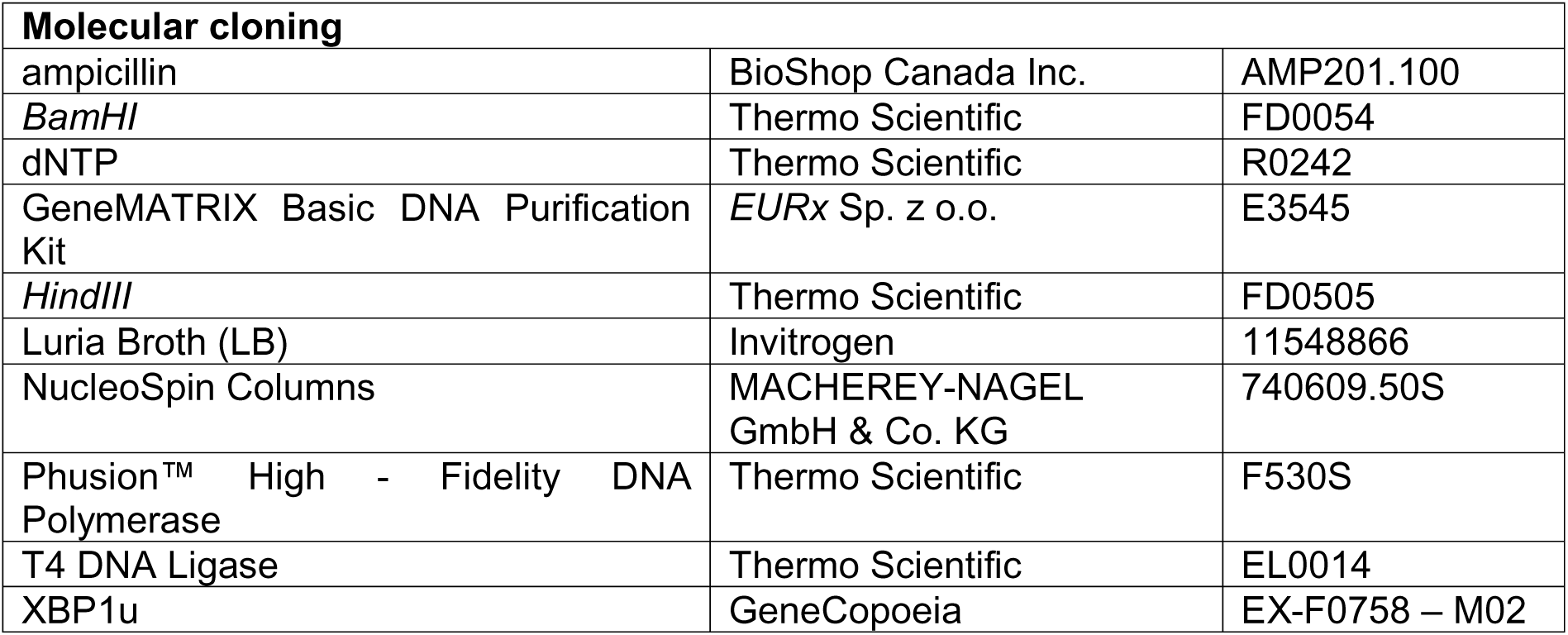

| **Molecular cloning** |  |  |
|---|---|---|
| ampicillin | BioShop Canada Inc. | AMP201.100 |
| *BamHI* | Thermo Scientific | FD0054 |
| dNTP | Thermo Scientific | R0242 |
| GeneMATRIX Basic DNA Purification Kit | *EURx* Sp. z o.o. | E3545 |
| *HindIII* | Thermo Scientific | FD0505 |
| Luria Broth (LB) | Invitrogen | 11548866 |
| NucleoSpin Columns | MACHEREY-NAGEL GmbH & Co. KG | 740609.50S |
| Phusion™ High - Fidelity DNA Polymerase | Thermo Scientific | F530S |
| T4 DNA Ligase | Thermo Scientific | EL0014 |
| XBP1u | GeneCopoeia | EX-F0758 – M02 |

### *in vitro* Transcription

The *XBP1u* transcript was generated by *in vitro* transcription from a *BamHI/HindIII*-linearized DNA template. After restriction digestion, the insert containing the *XBP1u* CDS was gel-purified using NucleoSpin columns. Transcription reactions were performed according to the manufacturer’s instructions using T7 Φ2.5 RNA polymerase (Thermo Scientific, EP0112) at a ratio of 50 units of enzyme per microgram of template DNA. ATP, GTP, CTP, and UTP (Thermo Scientific, R0441, R0461, R0451, and R0471, respectively) were added to reach a final concentration of 1 mM each. The reactions were then incubated at 37 °C for two hours in the presence of 20 U of RiboLock RNase inhibitor (Thermo Scientific, EO0381) and 5 mM of DTT (Sigma, 43815). The synthesized RNA was then purified using an RNA Clean & Concentrator-25 kit (Zymo Research, R1017), following the manufacturer’s protocol. The RNA concentration was determined spectrophotometrically using a Multiskan SkyHigh Microplate Spectrophotometer (Life Technologies, Warsaw, Poland) by measuring the absorbance at 260 nm (A₂₆₀).

| ***in vitro* transcription assay** |  |  |
|---|---|---|
| ATP | Thermo Scientific | R0441 |
| CTP | Thermo Scientific | R0451 |
| DTT | Sigma | 43815 |
| GTP | Thermo Scientific | R0461 |
| NucleoSpin Columns | MACHEREY-NAGEL GmbH & Co. KG | 740609.50S |
| RNA polymerase T7 Φ2.5 | Thermo Scientific | EP0112 |
| UTP | Thermo Scientific | R0471 |
| RiboLock RNase Inhibitor | Thermo Scientific | EO0381 |
| RNA Clean & Concentrator-25 columns | Zymo Research | R1017 |
| UTP | Thermo Scientific | R0471 |

### IRE1 cleavage assay

Purified *XBP1u* RNA was refolded by heating to 95°C followed by rapid cooling on ice for 2 min. Refolding buffer was then added to obtain final concentrations of 40 mM HEPES (pH 7.4), 50 mM KOAc and 10 mM Mg(OAc)₂. Samples were incubated at 37°C for 30 min to allow formation of RNA secondary structures. M2, control morpholino (ctrlM) or 4μ8C were preincubated separately with either recombinant IRE1α or refolded *XBP1u* RNA at a 1:1 molar ratio for 30 min on ice prior to initiation of the cleavage reaction.

Cleavage reactions were performed by incubating 0.5 μg recombinant human IRE1α kinase/RNase domain (amino acids 465–977; Sino Biological, 11905-H20B) with 8–10 μg refolded *XBP1u* RNA in reaction buffer containing 40 mM HEPES (pH 7.4), 50 mM KOAc, 10 mM Mg(OAc)₂, 5 mM DTT and 1 μM ADP for 30 min at 30°C, as previously described [15, 25]. Reactions were terminated by addition of an equal volume of 2× RNA Loading Dye (Thermo Scientific, R0641) followed by heat denaturation at 70°C.

Cleavage products were separated on 6% denaturing Novex™ TBE-Urea polyacrylamide gels (Invitrogen, EC6865BOX) in TBE running buffer. A RiboRuler Low Range RNA Ladder (Thermo Scientific, SM1831) was used as a molecular size marker. Gels were stained with SYBR Gold (Invitrogen, S11494), and RNA fragments were visualized using a ChemiDoc™ XRS imaging system (Bio-Rad).

| **IRE1 cleavage assay** |  |  |
|---|---|---|
| 2× RNA Loading Dye | Thermo Scientific | R0641 |
| IRE1α kinase/RNase domain | Sino Biological | 11905-H20B |
| Novex™ TBE-urea gels | Invitrogen | EC6865BOX |
| RiboRuler Low Range RNA Ladder | Thermo Scientific | SM1831 |
| SYBR Gold | Invitrogen | S11494 |

### Quantification and statistical analysis

All data are expressed as means ± SEM. For comparisons involving more than two groups, a one-way analysis of variance (ANOVA) was conducted, followed by Bonferroni’s post hoc test for multiple comparisons. Differences were considered significant at *p* < 0.05. All analyses were performed using GraphPad Prism version 10,6,0 (GraphPad Software, San Diego, CA). If representative results are presented, remaining results are available as datasets.

## Abbreviations

3′ UTR: 3′ untranslated region
4μ8C: IRE1 RNase inhibitor
ALLN: N-acetyl-Leu-Leu-norleucinal
ATF4: activating transcription factor 4
ATF6: activating transcription factor 6
BFA: brefeldin A
BiP: Heat Shock Protein Family A Member 5 (HSPA5)
BSA: bovine serum albumin
CHOP: DNA damage inducible transcript 3 (DDIT3)
DGAT2: diacylglycerol O-acyltransferase 2
DMSO: dimethyl sulfoxide
DTT: dithiothreitol
ER: endoplasmic reticulum
FBS: fetal bovine serum
GAPDH: glyceraldehyde 3-phosphate dehydrogenase
IRE1α: inositol-requiring enzyme 1 alpha
IXA4: IRE1 activator 4
MKC8866: IRE1 RNase inhibitor
PMO: phosphorodiamidate morpholino oligomer
RIDD: regulated IRE1-dependent decay
RT-qPCR: quantitative reverse transcription polymerase chain reaction
SERCA: sarco/endoplasmic reticulum Ca²⁺-ATPase
Thp: thapsigargin
Tm: tunicamycin
UPR: unfolded protein response;
XBP1: X-box binding protein 1
XBP1s: spliced X-box binding protein 1
XBP1u: unspliced X-box binding protein 1.

## DECLARATIONS

### Materials availability

Further information and requests for resources and reagents may be directed to and will be fulfilled by the corresponding authors.

### Data and code availability

All data reported in this paper will be shared by the lead contact upon request. This paper does not report original code. Any additional information required to reanalyze the data reported in this paper is available from the lead contact upon request.

## Acknowledgments

This work was funded by the National Science Centre, Poland (OPUS 2022/47/B/NZ7/00051 to R.B.), the Fondation pour la Recherche Médicale (FRM EQU202403018041 to E.C.), the Deutsche Forschungsgemeinschaft through SFB 1177 (Project F11; Project No. 259130777 to L.S.), and the Cardio-Pulmonary Institute (EXC 2026; Project ID 39064989 to L.S.). L.K was supported by Polish Ministry of Science and Higher Education, grant no. 4/SPUB/SP/0133/2026.

## Author contributions

Conceived and designed the experiments: A.W.W., R.B. E.C. Performed the experiments, A.W.W., M.G., J.S., D.P.R. Analyzed the data: R.B., A.W.W., M.G., J.S., E.C., D.P.R. Contributed reagents/materials/analysis tools: R.B., E.C., L.K., W.B., S.B. Wrote the manuscript: R.B., A.W.W. Revised the manuscript: L.S., J.F.C. and E.C.

## Declaration of generative AI and AI-assisted technologies

Generative AI language tools (Microsoft Copilot) were used to assist in correcting language and grammar in this document. No content was generated beyond linguistic refinement.

## FIGURE LEGENDS

**Supplementary Figure S1. Translation-blocking morpholino T1 does not efficiently suppress stress-induced XBP1s protein accumulation.** (**A**) Representative immunoblot analysis of XBP1s protein levels in HeLa cells transfected with control morpholino (ctrlM) or increasing concentrations of the translation-blocking morpholino T1 (0.5-2 μM), followed by tunicamycin (Tm) treatment. β-actin was used as a loading control. (**B**) Densitometric quantification of XBP1s protein levels shown in panel (A). Tm treatment induced robust accumulation of XBP1s protein, whereas T1 treatment did not produce a concentration-dependent reduction in XBP1s abundance. Data are presented as mean ± SEM. Statistical significance was determined by one-way ANOVA followed by Fisher’s LSD multiple-comparison test: ns, not significant; P < 0.05 (); P < 0.01 (**); P < 0.0001 (***). (**C**) Full, uncropped immunoblots together with TGX-stained membrane and gel images corresponding to the experiments shown in panel (**A**).

**Supplementary Figure S2. Additional splice-site-targeting morpholinos do not inhibit recombinant IRE1 kinase activity.** Recombinant IRE1 kinase activity was measured using an ADP-Glo kinase assay in the presence of increasing concentrations of the splice-site-targeting morpholinos M1, M3 and M4. None of the tested morpholinos produced a detectable reduction in IRE1 kinase activity across the concentration range analysed. Data are presented as mean ± SEM.

**Supplementary Figure S3. Functional heterogeneity of splice-site-targeting morpholinos.** Representative immunoblot analysis of XBP1s protein levels in HeLa cells exposed to tunicamycin (Tm) or thapsigargin (Thp) and treated with splice-site-targeting morpholinos M1– M4. Treatment with the selective IRE1 RNase inhibitor 4μ8C served as a positive control for inhibition of XBP1s production. Total protein staining is shown as a loading control. Although all morpholinos were designed to target the XBP1 splice substrate, substantial differences in inhibitory activity were observed, with M2 consistently producing the strongest reduction in stress-induced XBP1s accumulation.

**Supplementary Figure S4. Full-length Phos-tag immunoblots corresponding to Figure 6A**. Full-length Phos-tag SDS-PAGE immunoblots of total and phosphorylated IRE1 (P-IRE1) in HeLa cells transfected with control morpholino (ctrlM) or M2 and exposed to tunicamycin (Tm) or thapsigargin (Thp). Cells treated with the IRE1 RNase inhibitor 4μ8C were included as a control. Corresponding β-actin immunoblots are shown below. The cropped regions used for preparation of **Figure 5A** are indicated in the lower panels.

**Supplementary Figure S5. Full immunoblot dataset used for quantification of XBP1s protein levels in Figure 2D**. Representative full-length immunoblots from five independent biological replicates used for quantification of XBP1s protein abundance presented in **Figure 2D**. HeLa cells were treated with tunicamycin (Tm) or thapsigargin (Thp) in the presence or absence of candidate morpholinos and the IRE1 inhibitor 4μ8C. These data are provided to document the reproducibility of XBP1s protein measurements and the source blots used for densitometric analysis.

