## Supplemental Figures for "Targeting the *XBP1* splice substrate redirects IRE1-mediated RNA processing without direct inhibition of IRE1"

**Figure S1**

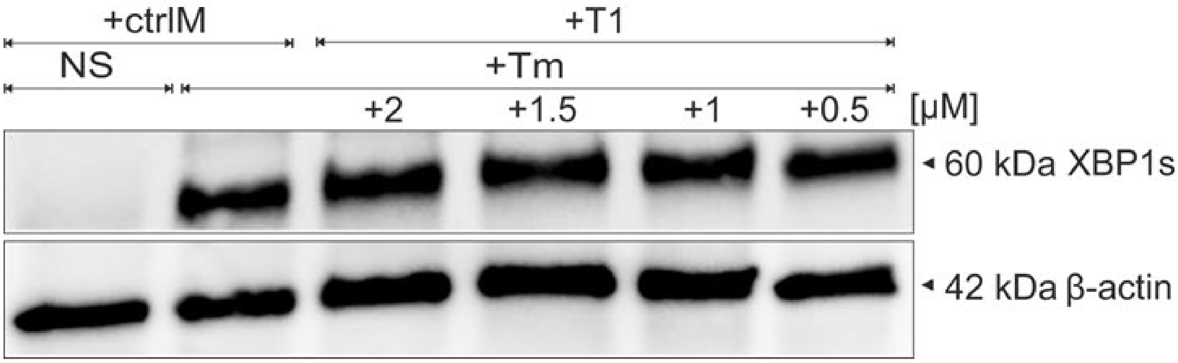
+ctrlM
+T1
NS
+Tm
+2
+1.5
+1
+0.5
[µM]
60 kDa XBP1s
42 kDa β-actin

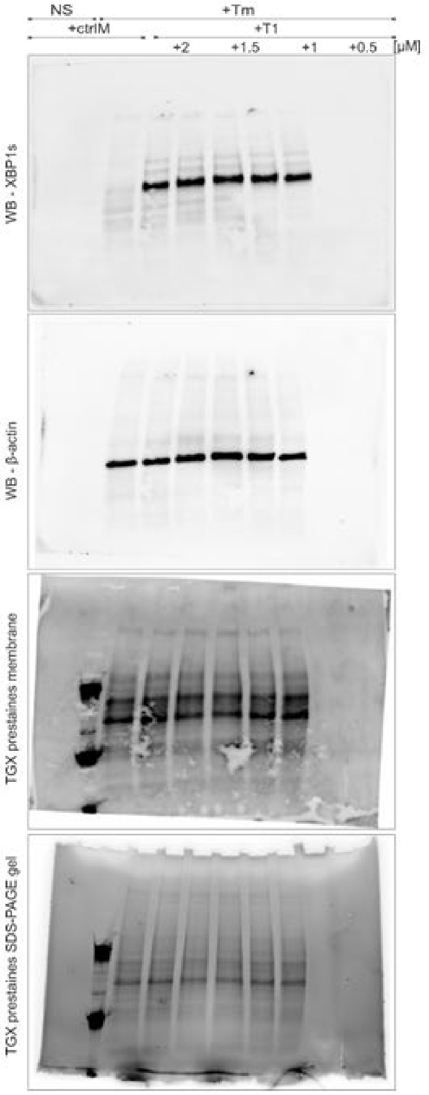
NS
+Tm
+ctrlM
+T1
+2
+1.5
+1
+0.5
[µM]
WB - XBP1s
WB - β-actin
TGX prestaines membrane
TGX prestaines SDS-PAGE gel

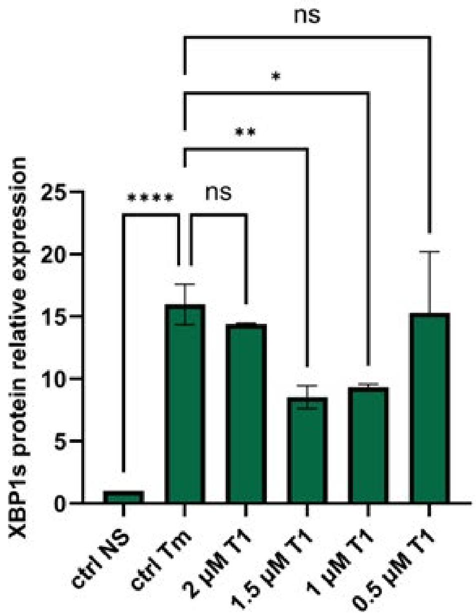
ns
*
**
****
ns
XBP1s protein relative expression
25
20
15
10
5
0
ctrl NS
ctrl Tm
2 µM T1
1.5 µM T1
1 µM T1
0.5 µM T1

**Figure S2**

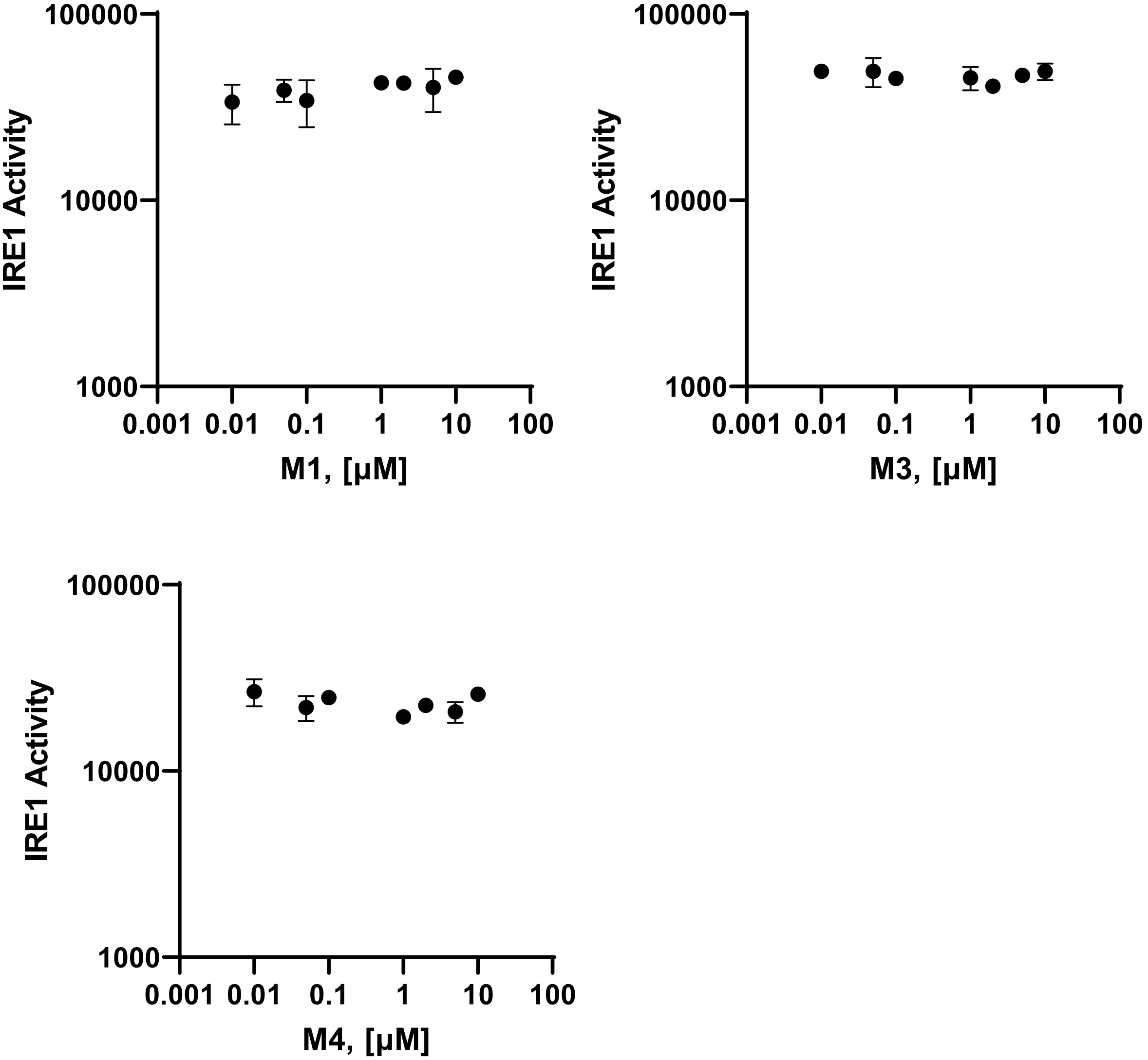
IRE1 Activity
100000
10000
1000
0.001 0.01 0.1 1 10 100
M1, [µM]
IRE1 Activity
100000
10000
1000
0.001 0.01 0.1 1 10 100
M3, [µM]
IRE1 Activity
100000
10000
1000
0.001 0.01 0.1 1 10 100
M4, [µM]

**Figure S3**

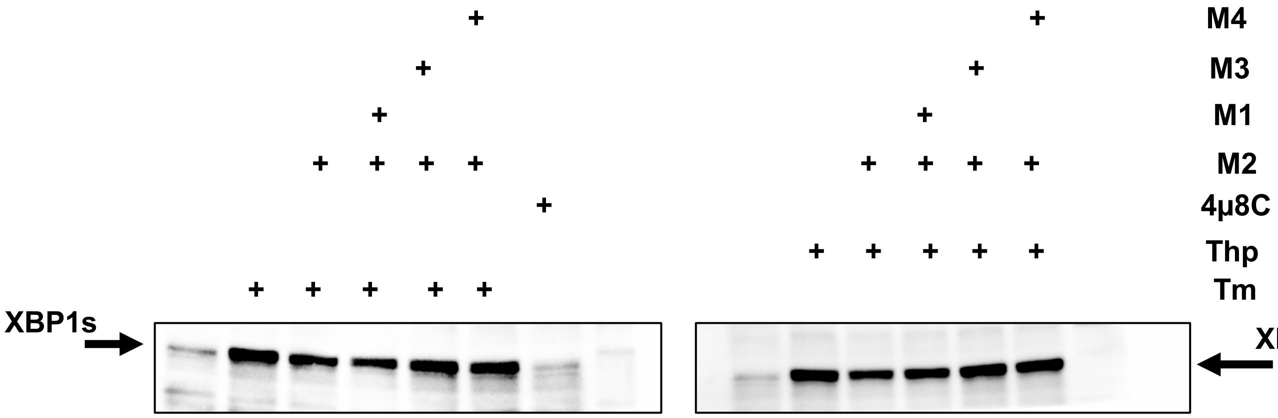


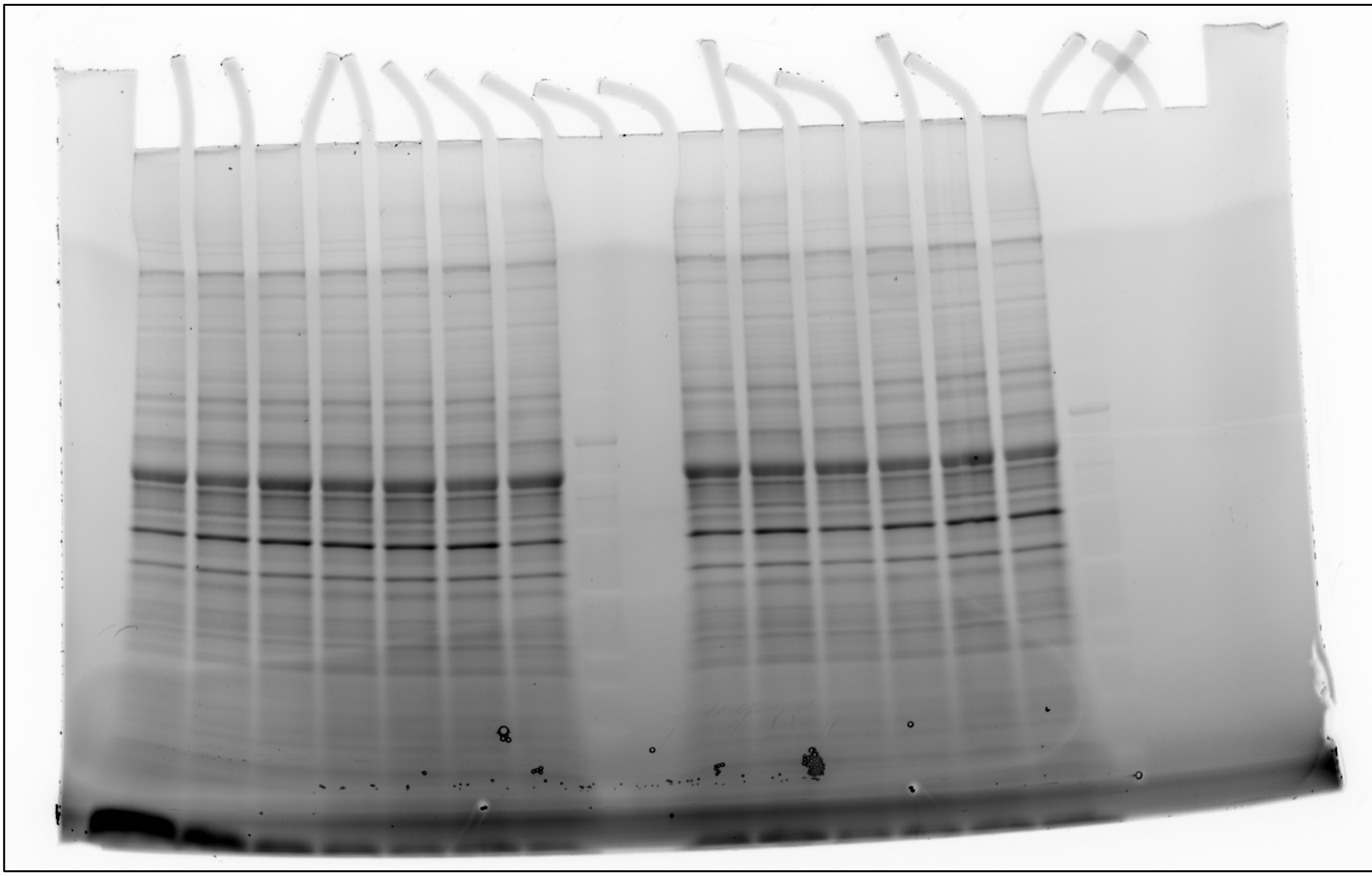

**Total protein**

### Figure S4

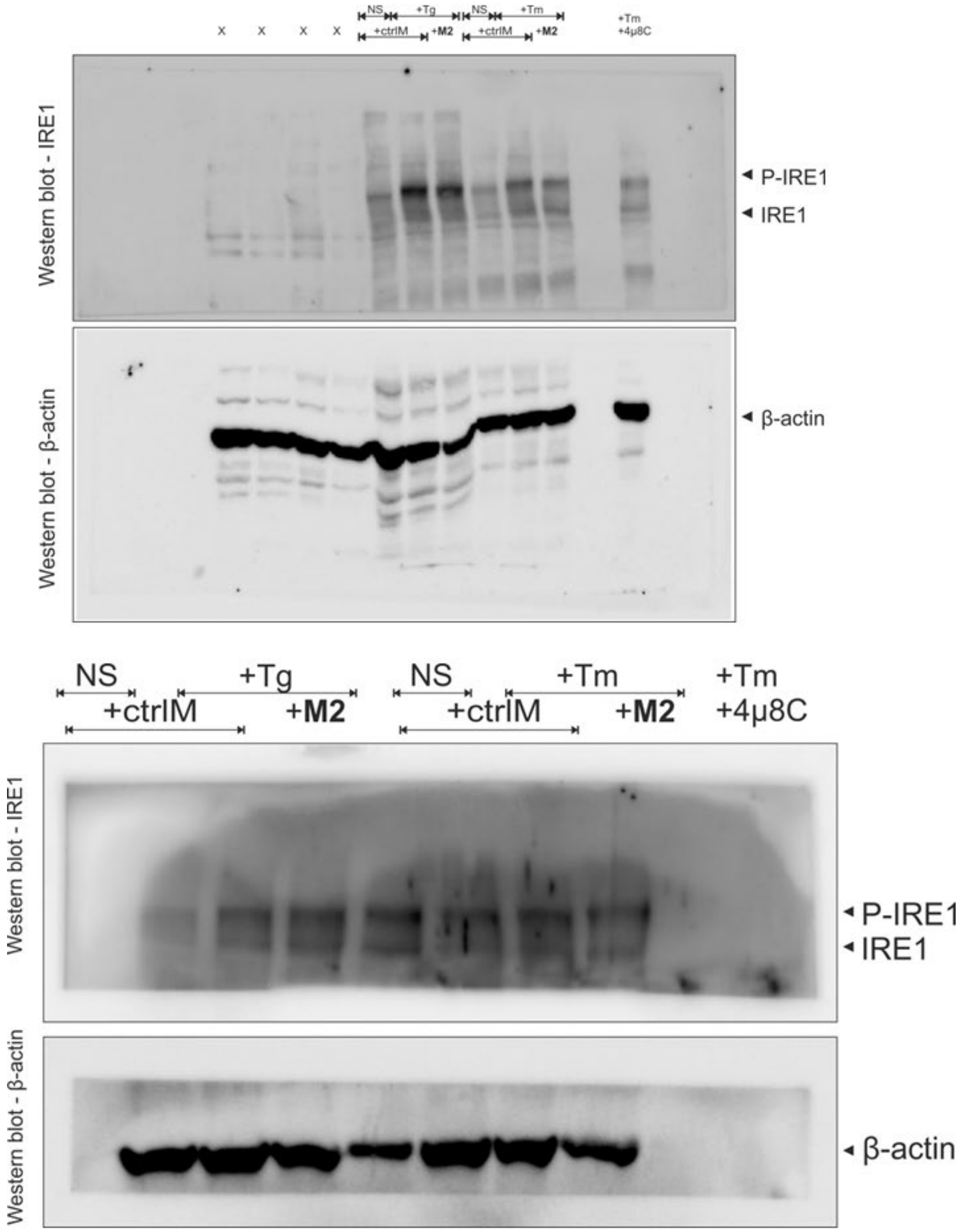
x x x x
NS
+Tg
NS
+Tm
+ctrlM
+M2
+ctrlM
+M2
+Tm
+4μ8C
Western blot - IRE1
P-IRE1
IRE1
Western blot - β-actin
β-actin
NS
+Tg
NS
+Tm
+Tm
+ctrlM
+M2
+ctrlM
+M2
+4μ8C
Western blot - IRE1
P-IRE1
IRE1
Western blot - β-actin
β-actin

#### Figure 2 D.

##### Replicate 1

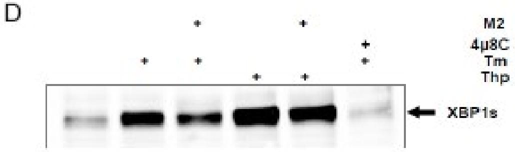


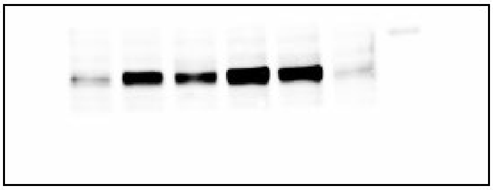

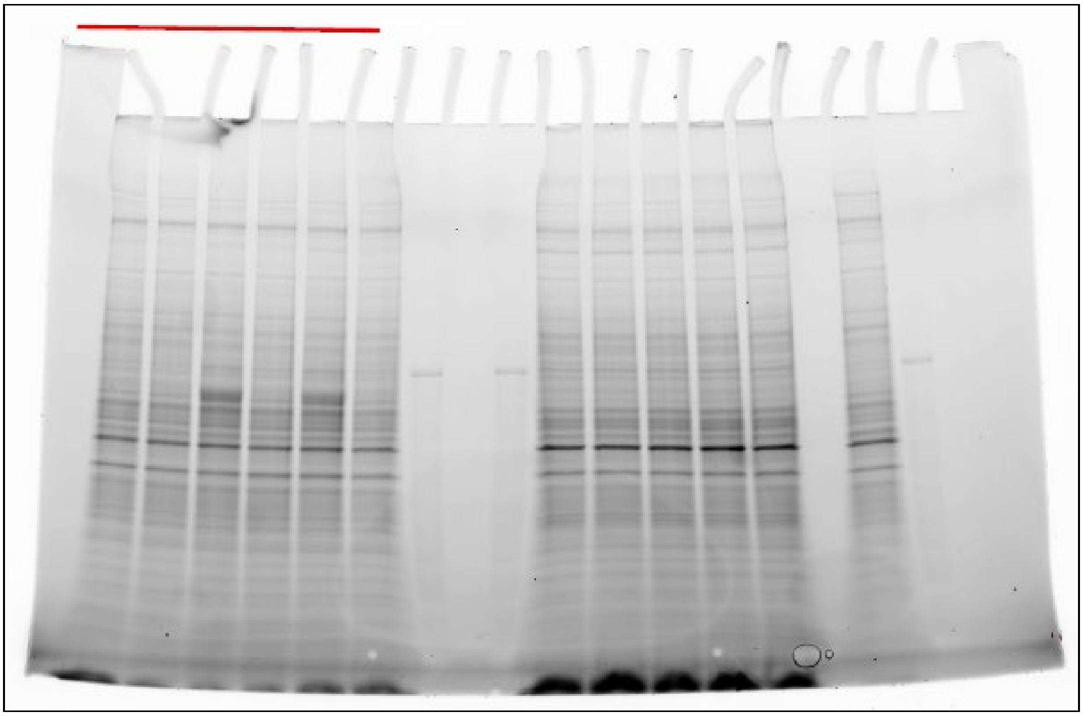

**Figure S5**

**Replicate 2 (same sample order)**

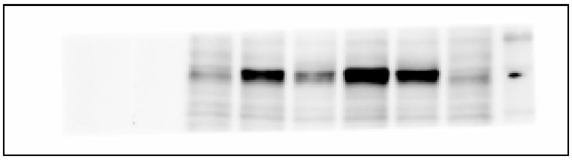

**Replicate 3 (same sample order)**

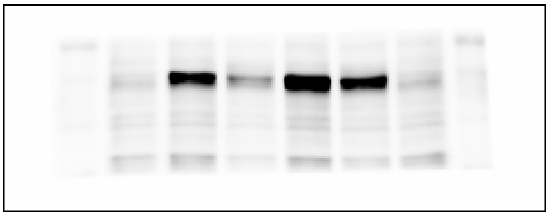

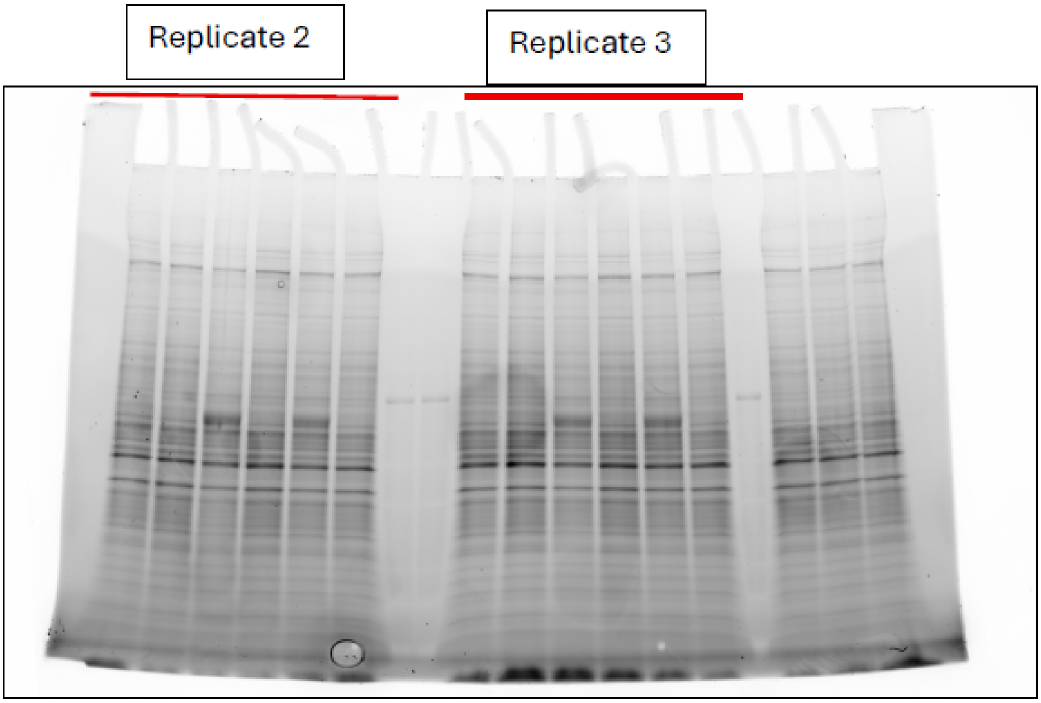


**Replicate 4**

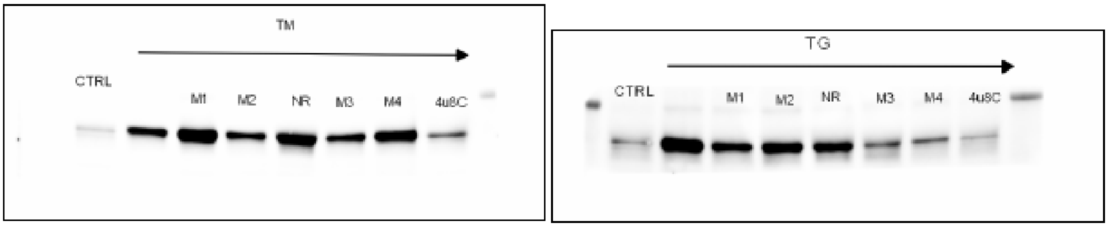

**Figure S5**

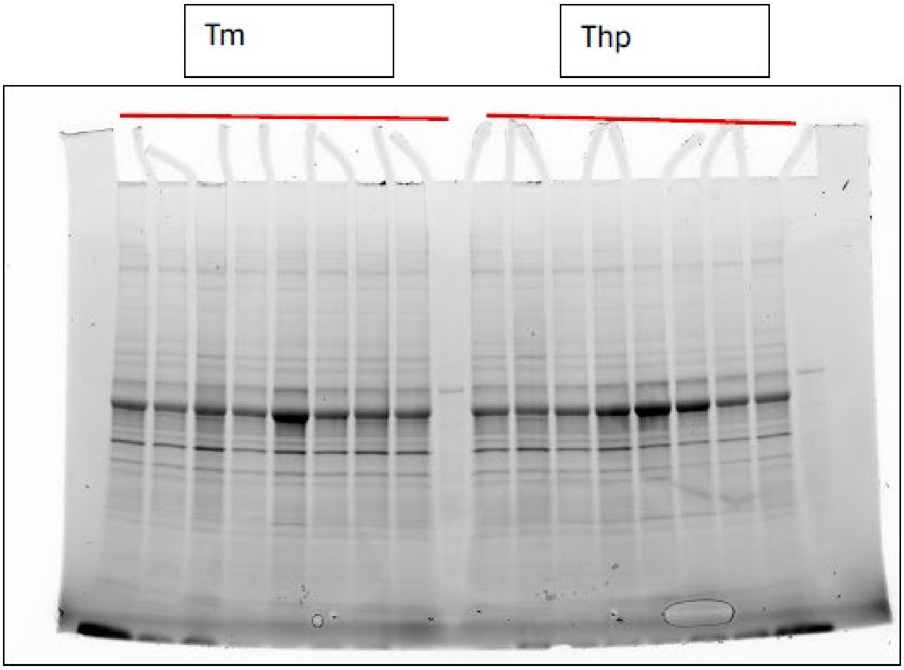


**Replicate 5**

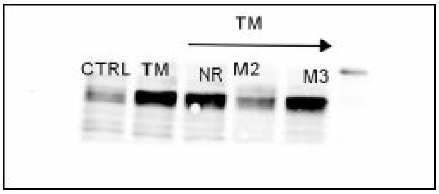


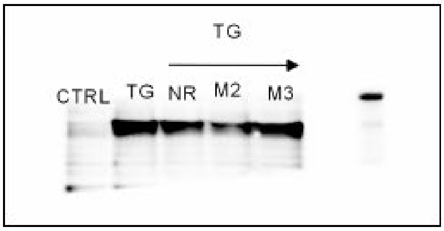


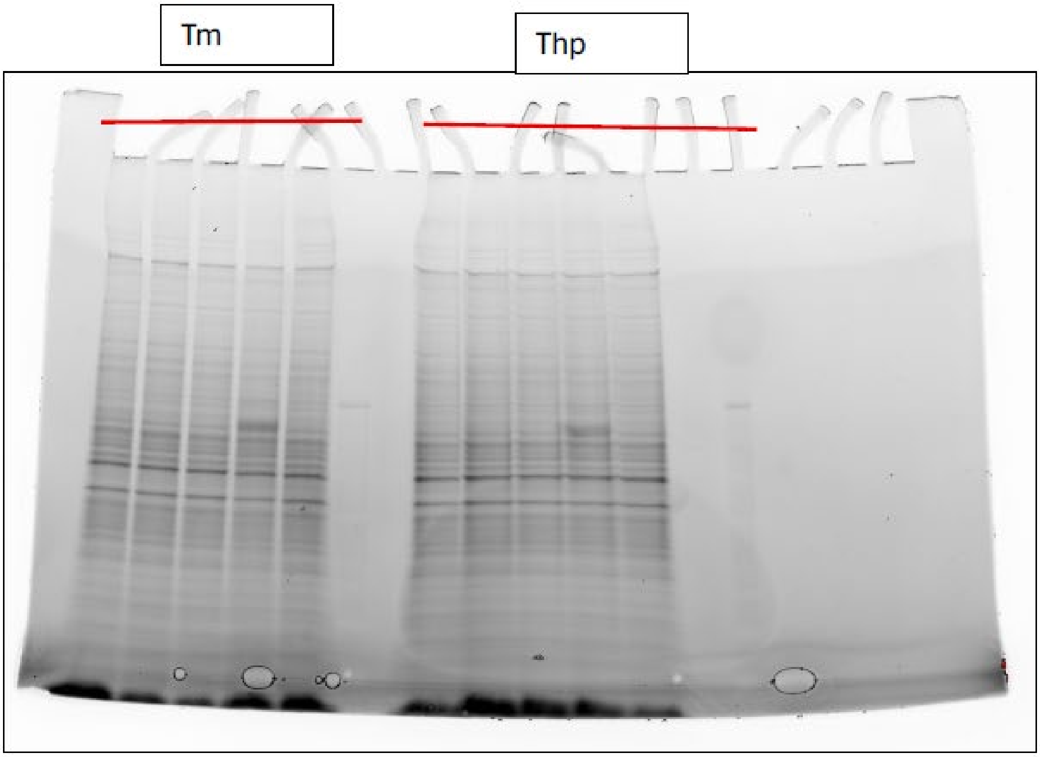
